# Interspecific hybrids reveal a deeply conserved nuclear role of EIF1A-family proteins in zygotic genome activation

**DOI:** 10.64898/2026.10.26.761752

**Authors:** Ding Qiu, Dengfeng Bi, Zhiheng Yuan, Zhenghao Qiao, Zhongcheng Zhou, Zeling Wang, Yunlong Li, Zhenyu Zhong, Yibei Wu, Ziyan Rong, Lijun Deng, Xiaohua Lu, Tong Wang, Yongkang Jia, Jiemei Chen, Weiwei Liu, Sheng Wang, Xiaozhong Zhang, Tianyu Zhang, Ming Wang, Quanyong Zhang, Xiaoyun Wang, Wenhua Yu, Yanna Dang, Chao Liu, Wei Li, Zhiheng Chen, Ling Sun, Siqi Gui, Jingcheng Zhang, Jie Li, Yunkun Dang, Yuan Qin, Tiejun Zhang, Hao Chen, Qianqian Sha, Yanfang Wang, Na Yang, Wei Xie, Kai Chen, Jianguo Zhao, Yang Yu

## Abstract

Zygotic genome activation (ZGA) launches inaugural gene expression and autonomous development of the embryo. Yet, conserved regulators of ZGA among different mammals remain poorly defined. Here, we generated interspecific hybrid embryos to place distant mammalian genomes in a shared maternal cytoplasm. These hybrids undergo cleavage and transcriptional activation from divergent parental genomes. Comparative transcriptomics identifies a core set of zygotic genes reproducibly activated across different hybrid contexts, including the *Eif1a-like* family. Depletion of EIF1A-like proteins disrupts ZGA and preimplantation development in mouse, pig and cattle embryos. Although best known for translation initiation, these proteins contain a conserved nuclear localization signal and translocate to the nucleus at the onset of major ZGA. Further studies revealed that they occupy promoters of nearly all ZGA genes and MERVL/MT2 retrotransposons, required for local chromatin accessibility. These findings uncover a conserved nuclear licensing function for EIF1A-family proteins in zygotic genome activation among divergent mammalian embryos.

## Main Text

At fertilization, the embryo inherits abundant maternal RNAs and proteins but a largely silent genome. Development becomes autonomous only when the zygotic genome is activated, unleashing transcription of thousands of genes and retrotransposons(*1*). The transition of a fertilized egg from maternal control to its own developmental program represents a universal transition in nearly all metazoan(*2, 3*). Yet, its timing and transcriptional output vary substantially among mammals(*4, 5*). Whether such diversity is built on conserved principles or shared molecular machinery remains unresolved.

At the molecular level, ZGA is orchestrated by regulators that selectively recognize defined genomic loci to coordinate chromatin accessibility, epigenetic remodeling, and the recruitment of the transcriptional apparatus(*1, 2, 6*). While most knowledge comes from species-specific studies, several key ZGA-specifying factors, like DUX and OBOX/TPRX families, have analogous roles in mouse and human, but with limited conservations(*7–14*). However, it remains unclear whether deeper conserved principles govern ZGA across mammals, largely due to the lack of systematic genetic perturbations to interrogate ZGA requirements(*2, 5, 15, 16*). Moreover, conventional cross-species comparisons inherently confound the contributions of the oocyte cytoplasm and the embryonic genome, potentially obscuring conserved logic(*17*). If such general principles exit, the regulatory machinery of one species would be sufficient to activate the genome of another. To answer this question, we placed divergent genomes into a foreign cytoplasm by intracytoplasmic injecting sperm (ICSI) of one species into oocytes of another(*18*). If these hybrids support cell cleavage and transcriptional activation, they would enable us to uncover general principles governing ZGA across mammals. Genes that are reproducibly activated across hybrid contexts are therefore highly correlated with successful ZGA, which would represent strong candidates for maintaining conserved ZGA functions.

Here, we profile hybrid embryos from mammals separated by over 60 million years of evolution. We demonstrate that divergent genomes can undergo extensive ZGA in a foreign oocyte cytoplasm. Comparative analysis identifies EIF1A-like protein family as core ZGA licensing factors. These proteins relocalize to the nucleus during ZGA to maintain chromatin accessibility at ZGA promoters and retrotransposons, uncovering a noncanonical nuclear function for a classical translation factor in early embryogenesis.

### Interspecies hybrid embryos undergo early cleavages

Mammalian species differ substantially in both the timing and transcriptional programs of ZGA(*4*) (Fig. 1A and fig. S1A). To test whether mammalian ZGA is governed by conserved regulatory principles, we asked whether a foreign oocyte could activate an evolutionarily distant sperm genome (fig. S1B). We therefore generated interspecies hybrid embryos by intracytoplasmic sperm injection (ICSI) using oocytes from mouse or pig and sperm from mouse, pig or cattle. Such hybrids have been produced for decades, yet their transcriptomes remained unknown (fig. S1C)(*19–21*). We established four hybrid combinations: porcine oocyte-mouse sperm (PM), porcine oocyte-bovine sperm (PB), mouse oocyte-porcine sperm (MP) and mouse oocyte-bovine sperm (MB) (Fig. 1B). Most hybrid embryos arrested at the 2-cell stage, but a subset continued cleaving, reaching the 8-cell stage and thereby spanning the normal ZGA windows of both parental species (Fig. 1C and fig. S1D). For example, in MP embryos, nuclear staining resolved distinct maternal and paternal pronuclei, the latter forming by 9 hours post-ICSI, slower than in conspecific controls (6 hours) (fig. S1E). Cleavage nevertheless proceeded: by 24 and 48 hours post-ICSI, these embryos had reached the two-cell and four-cell stages, respectively. Thus, early cleavage can proceed even when an embryo carries genomes from two distantly related mammals.

**Fig. 1.**
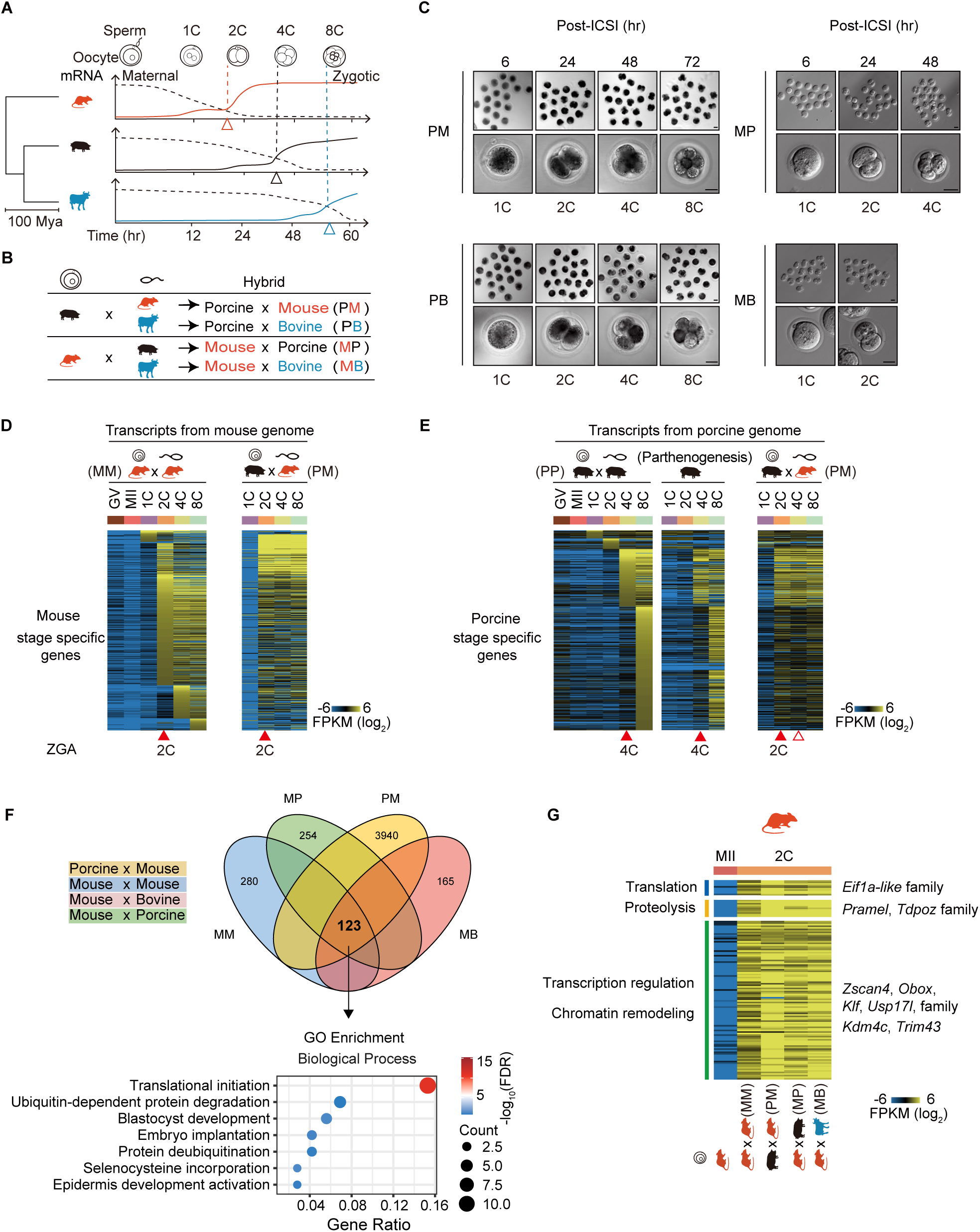
Interspecific hybrid embryos reveal conserved ZGA programs. (**A**) Schematic of maternal-to-zygotic transition timing in mouse, pig and cattle. Arrowheads mark the onset of major ZGA. (**B**) Oocyte–sperm combinations used to generate interspecies hybrid embryos. (**C**) Bright-field images of hybrid embryos at the indicated hours post-ICSI, with the cleavage stages reached. Scale bars, 20 μm. (**D**) Expression of mouse stage-specific genes in natural (MM) and hybrid (PM) embryos. (**E**) Expression of porcine stage-specific genes in natural (PP), parthenogenetic and hybrid (PM) embryos. In d and e, filled arrowheads mark the observed onset of ZGA; the open arrowhead in e marks the stage at which the porcine genome is normally activated (4C). (**F**) Overlap of genes activated in MM, PM, MP and MB embryos and Gene Ontology enrichment of the 123 shared genes. (**G**) Expression of the shared genes at MII and two-cell stages across all four combinations, grouped by function.

### Divergent genome zygotic activation in hybrid embryos

We next asked whether the zygotic genomes in these hybrids undergo transcriptional activation. 5-Ethynyl uridine (EU) labeling revealed robust nascent RNA synthesis in 2-cell MP embryos, comparable to that in normal mouse embryos (fig. S1F). We therefore performed single-embryo SMART-seq on morphologically normal hybrids collected at stages matched to normal development (e.g., 2-cell embryos at 24 hours and 4-cell embryos at 48 hours post-ICSI)(*22–24*). Because the parental species diverged more than 60 million years ago, most reads could be assigned unambiguously to the maternal or paternal genome. As sperm contributes negligible RNA to the zygote(*25*), reads assigned to the paternal genome provided a direct readout of zygotic transcription.

We sequenced 20-30 embryos per stage (1-, 2-, 4-, and 8-cell whenever available) for each hybrid combination. Nearly one-third of 2-cell PM embryos contained abundant transcripts derived exclusively from the paternal mouse genome (fig. S2A). On average, ∼27% of SMART-seq reads mapped unambiguously to the mouse genome, indicating extensive paternal genome activation. This fraction is likely an underestimate given existing maternal RNAs. Notably, the paternal mouse genes activated in PM hybrids closely matched those induced in normal mouse embryos (Fig. 1D), indicating that porcine ooplasm can execute a broadly mouse-like ZGA program. This suggests that mouse and porcine may share conserved mechanisms for activating the paternal genome.

The PM hybrids also altered the maternal porcine ZGA program. Abundant porcine transcripts appeared at the 2-cell stage together with paternal mouse transcripts (Fig. 1E, right panel). To exclude maternal deposited RNA, only transcripts with counts exceeding RNA levels of metaphase II (MII) oocyte were considered as confident zygotic expression(*15, 26, 27*). Notably, the maternal porcine genome was activated simultaneously with the paternal mouse genome at the 2-cell stage in PM hybrids. This contrasts sharply with normal or parthenogenetic porcine embryos, in which major ZGA occurs at the 4-cell stage (Fig. 1E, left panels). Analogously, both parental genomes were likewise transcriptionally active at the 2-cell stage in reciprocal MP hybrids (fig. S2, B to D). Like ZGA genes, mouse LTR retrotransposons were induced in these hybrids(*28, 29*) (fig. S2, E and F).

Similarly, Bovine sperm activate ZGA with maternal genomes, at the 4-cell stage in Porcine oocyte and 2-cell stage in mouse oocytes respectively (fig. S2G). Thus, a shared embryonic environment allows synchronization of normally distinct ZGA programs. Across all crosses, foreign oocytes activated divergent sperm genomes, suggesting that the regulatory principles underlying ZGA are deeply conserved across mammals. Notably, every hybrid containing a mouse genome initiated ZGA at the 2-cell stage, irrespective of cross direction (fig. S2G). Remarkably, the porcine genome in PM embryos was activated one cleavage cycle ahead of its normal schedule. These results indicate that mouse-encoded regulatory information is sufficient to drive ZGA in porcine and bovine genomes, even in a foreign embryonic context.

Hybrid embryos also exposed a normally hidden component of ZGA. Housekeeping transcripts are abundant maternal products and are therefore difficult to distinguish from newly synthesized RNA in conventional embryos, although their zygotic activation has been detected in fish and sea squirt hybrids(*30–32*). In PM hybrids, ∼64% of mouse housekeeping genes (708 of 1,103) were transcribed from the paternal genome (fig. S2H); in MP hybrids, ∼79% of porcine housekeeping transcripts (467 of 594) were transcribed from the paternal genome (fig. S2I). Housekeeping genes may therefore constitute a substantial, previously underappreciated component of mammalian ZGA, potentially universal across metazoans.

### Hybrid embryos nominate a family of *Eif1a-like* genes as candidate ZGA regulators across species

The precocious activation of porcine genes in PM embryos suggested that paternal mouse genome produces factors capable of advancing the host program. We therefore searched for mouse genes consistently induced across hybrids carrying a mouse genome, identifying 123 such genes activated at the 2-cell stage in all contexts (Fig. 1F). Gene Ontology analysis highlighted “Translation” and “Proteolysis”, alongside established ZGA regulators, including members of *Obox*, *Zscan4*, and *Usp17l* families, as well as *Zfp352*, *Kdm4c*, and *Klf9*(*12, 13, 33–36*) (Fig. 1G and fig. S3A). The GO term “Translation” was dominated by a family of highly similar *Eif1a-like* genes clustered on chromosome 12, together with *Eif1a* on chromosome 18(*37, 38*) (Fig. 2A). These genes were transcribed and translated specifically at the 2-cell stage in both normal mouse and hybrid embryos(*39*) (Fig. 2B and fig. S3, B and C). Additional *Eif1a* family members, like *Eif1ax* on the X chromosome, showed a similar activation pattern(*40, 41*) (fig. S4A).

**Fig. 2.**
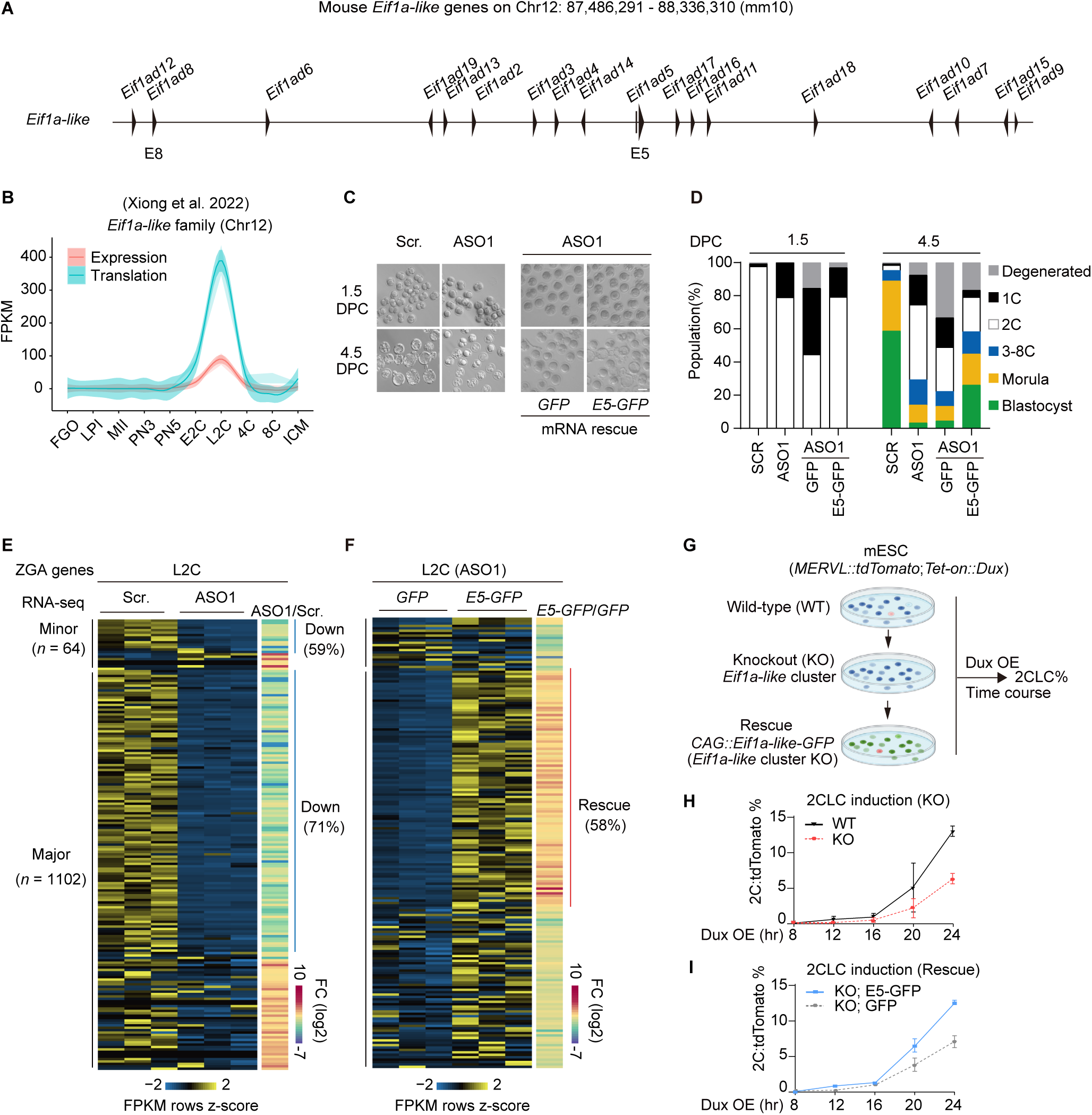
The *Eif1a-like* cluster is required for mouse ZGA. (**A**) Genomic organization of the mouse *Eif1a*-*like* gene cluster on chromosome 12 (mm10). The members used for rescue constructs (E5, E8) are indicated. (**B**) Expression (RNA) and translation (RPF) dynamics of *Eif1a-like* family genes across mouse oocyte and preimplantation stage. (**C**) Representative bright-field images of scrambled-control (Scr.), ASO1-injected and mRNA-rescued embryos at 1.5 and 4.5 days post coitum (DPC). Scale bar, 100 μm. (**D**) Developmental stage distribution of embryos in c at 1.5 and 4.5 DPC. (**E**) Expression of minor (*n* = 64) and major (*n* = 1,102) ZGA genes in Scr. and ASO1-injected late two-cell (L2C) embryos, with ASO1/Scr. fold changes. (**F**) Expression of the same gene sets in ASO1-injected embryos rescued with *GFP* or *E5-GFP* mRNA. (**G**) Schematic of the 2CLC induction assay in wild-type (WT), *Eif1a*-*like* cluster-knockout (KO) and E5-GFP-rescued mESCs carrying *MERVL::tdTomato* and *Tet-on::Dux* reporters. (**H** and **I**) 2C::tdTomato-positive cells after Dux overexpression in WT versus KO mESCs (H) and in KO mESCs rescued with GFP or E5-GFP (I). Data are mean ± s.e.m. of n = 3 independent inductions;

### *Eif1a-like* genes are conserved in sequence and zygotic expression pattern

Phylogenetic analysis showed a lineage-specific expansion of the *Eif1a-like* subfamily specifically in mouse, whereas the pig, cattle and human genomes each contained *Eif1ax* and/or *Eif1ay* homologs (fig. S4B). These homologs were also induced during species-specific ZGA (fig. S4C). Interestingly, human EIF1A homologs were strongly induced at the 8-cell stage (fig. S4D) and followed normal induction timing in human androgenetic embryos (paternal genome only), which maintained correct ZGA timing(*42*). In contrast, they are reduced in parthenogenetic embryos (maternal genome only), in which ZGA is delayed(*43*) (fig. S4E). Across multiple mammalian species, EIF1A, EIF1AX/Y, and representative EIF1A-like proteins (mouse EIF1AD5 and EIF1AD8) all shared a conserved oligonucleotide-binding (OB) fold domain flanked by unstructured N- and C-terminal tails(*41*) (fig. S4F). For simplicity, we collectively refer to all *Eif1a* members as EIF1A-family proteins, including EIF1A, EIF1AX/Y and EIF1A-like (EIF1AD2-19).

### The EIF1A-like family is required for ZGA across mammals

To test whether the *Eif1a-*family is functionally required for mouse preimplantation development, we designed two antisense oligonucleotides against an open reading frame region shared by all family members, enabling simultaneous depletion of all *Eif1a-like* transcripts (fig. S5A). Both ASOs arrested early embryonic development relative to the scramble control (Fig. 2C and D and fig. S5, B to D). Single-embryo RNA-seq of late 2-cell embryos (L2C) injected with ASO1 revealed broad collapse of the ZGA program: 59% of minor ZGA genes (38 of 64) and 71% of major ZGA genes (778 of 1,102) were downregulated (Fig. 2E and fig. S5E). ASO2 produced a similar effect (fig. S5, F and G), and the suppressed genes substantially overlapped with those altered in maternal-zygotic *Obox-*family knockout embryos(*12*) (fig. S5H). An ASO-resistant *Eif1ad5-GFP* mRNA (*E5-GFP*), but not GFP alone, partially restored development and ZGA-gene expression (Fig. 2, C, D and F). Depletion of *Eif1ax* genes in porcine and bovine embryos caused a similar arrest (fig. S5, I to L). EIF1A*-*like protein activity is therefore conserved and required for ZGA and early development across the mammals tested.

### The Eif1a-like cluster is required for 2-cell-like cell conversion in mESCs

The expression of the Eif1a-like cluster was similarly coupled to the 2-cell program in mouse embryonic stem cells (mESCs) (fig. S6A). They were induced in spontaneously appearing 2-cell-like cells (2CLCs) and after DUX or OBOX5 overexpression(*7, 12, 44*) (fig. S6, A to D). Using CRISPR/Cas9, we deleted the entire *Eif1a-like* cluster (∼0.7Mb) on chromosome 12 in mESCs carrying a MERVL::tdTomato 2C reporter(*29, 45*) (Fig. 2G and fig. S6, E and F). Cluster deletion sharply reduced DUX-induced 2CLC conversion (Fig. 2H). Overexpression of E5-GFP, but not GFP alone, restored the response, as did an EIF1AD8-GFP fusion (E8-GFP) in two independent clones (Fig. 2I and fig. S6G). The Eif1a-like cluster is therefore important for the 2CLC transition, mirroring its role in embryonic ZGA.

### Nuclear EIF1A-like proteins are required for ZGA

EIF1A-like proteins are best known as components of the translation-initiation machinery(*40*), prompting us to examine where they act during ZGA. In stark contrast to their presumed role in cytoplasm, antibody staining revealed strong nuclear accumulation in 2-cell mouse embryos(*46*) (Fig. 3A, left panel and fig. S6H). In porcine and bovine embryos, nuclear signal instead peaked at the 4- and 8-cell stages, respectively, matching the species-specific timing of major ZGA(*26, 47*) (Fig. 3A). E5-GFP and E8-GFP mRNA injection reproduced the nuclear pattern in mouse 2-cell embryos, whereas GFP was diffusely distributed throughout the embryo (Fig. 3B). Sequence comparison revealed a highly conserved N-terminal nuclear localization signal (NLS) that is invariant from mammals to nematodes, even as the rest of the protein, including the central OB-fold domain, accumulates substitutions (Fig. 3C). The coincidence of nuclear entry with ZGA across species suggests an evolutionarily conserved nuclear function for EIF1A-like proteins, beyond their canonical role in cytoplasmic translation.

**Fig. 3.**
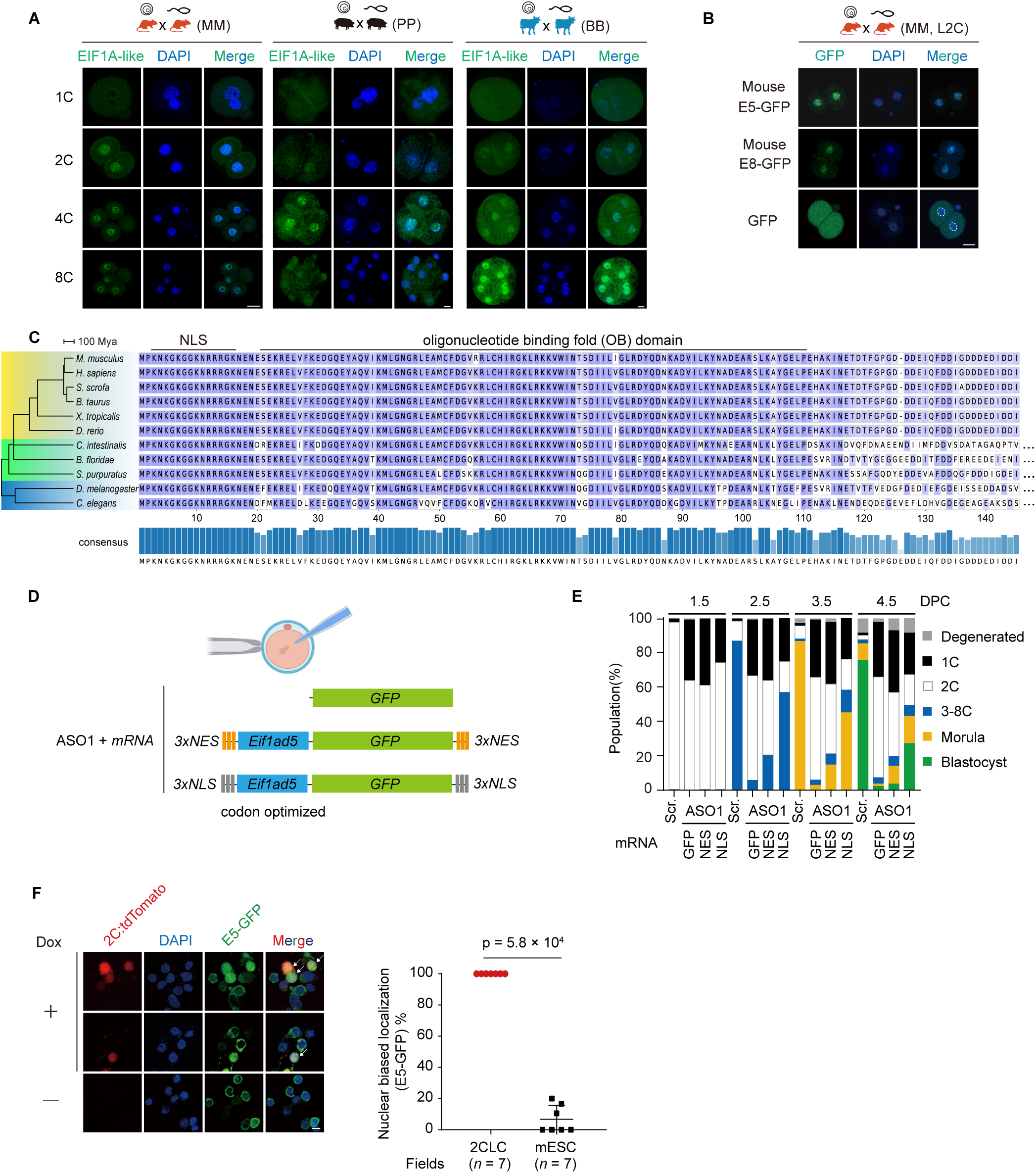
A deeply conserved nuclear function for EIF1A-like proteins during ZGA across mammals. (**A**) Immunofluorescence of EIF1A-like proteins in mouse (MM), pig (PP) and bovine (BB) embryos at the indicated stages. DNA was counterstained with DAPI. Scale bar, 20 μm. (**B**) Localization of E5-GFP (Eif1ad5–GFP), E8-GFP (Eif1ad8–GFP) or GFP alone in late two-cell mouse embryos. Scale bar, 20 μm. (**C**) Amino acid sequence alignment of EIF1A-family proteins across species spanning million years of evolution. The nuclear localization signal (NLS) and oligonucleotide-binding (OB) domain are indicated; the consensus sequence is shown below. (**D**) Schematic of the rescue experiment: ASO1 injection followed by codon-optimized, ASO1-resistant *Eif1ad5* mRNA fused to GFP and flanked by triple NLS (3xNLS) or nuclear export signals (3xNES). (**E**) Developmental stage distribution of embryos at the indicated days post coitum after the treatments in d. (**F**) Nuclear-biased localization of E5-GFP in 2CLCs and mESCs. Each data point represents the percentage of cells with predominantly nuclear E5-GFP signal in one field of view (n = 7 fields per condition, from two independent induction experiments; 31 cells for 2CLC and 51 cells for mESC in total). P = 5.8 × 10⁻⁴, two-tailed Mann–Whitney U test.

To test whether nuclear localization is functionally required for ZGA, we depleted endogenous *Eif1a-like* transcripts with ASO1 and co-injected ASO-resistant E5-GFP fused with six tandem NLS or nuclear exporting signal (NES) motifs(*48, 49*) (Fig. 3D). ASO1 alone arrested most embryos at the 2-cell stage. NLS-E5-GFP, partially rescued development, whereas NES-E5-GFP and GFP did not (Fig. 3E). Therefore, precise nuclear localization is required for EIF1A-like function during ZGA. We then asked whether nuclear localization is also coupled to the 2-cell like state. We engineered mESCs carrying the MERVL::tdTomato 2C reporter to stably express E5-GFP(*45*). Dux overexpression could drive these mESCs into a 2CLC state marked by tdTomato(*29*). Strikingly, E5-GFP accumulated in the nuclei of these 2CLCs but remained largely cytoplasmic in tdTomato-negative or uninduced mESCs (Fig. 3F). Thus, EIF1A-like proteins relocalize into nucleus only when cells enter a 2C-like state or undergo ZGA.

### EIF1A-like proteins occupy ZGA gene promoters

Because DUX-induced 2CLCs recapitulated EIF1A-like nuclear localization, we used this system to investigate how these proteins act in a 2C-like state. We performed chromatin-immunoprecipitation sequencing (ChIP-seq) and CUT&Tag on mESCs stably expressing E5-GFP, before and after DUX-induced transition to 2CLCs (Fig. 4A). ChIP-seq of E5-GFP identified 21,580 peaks in 2CLCs that were largely absent from mESCs (Fig. 4B), consistent with 2CLC-specific nuclear accumulation. About 55% of peaks mapped to genes; the remaining distal peaks were dominated by transposons (∼76%), including LTRs (45%), LINEs (28%), and SINEs (3%) (Fig. 4C). Two occupancy patterns emerged: narrow peaks at GC-rich regions and broad peaks across MERVL retrotransposons (Fig. 4D). Gene-associated peaks centered on transcription start sites (TSSs), whereas transposon-associated signal followed their transcribed regions (fig. S7, A to D). E5-GFP was particularly enriched at MT2, the MERVL LTR that contains the retrotransposon promoters, producing strong signal at both ends of full-length MERVL elements (fig. S7, E and F). By contrast, 2CLC enhancers marked by H3K4me1 and H3K27ac showed little E5-GFP enrichment despite being accessible(*50*) (fig. S7G). Thus, EIF1A-like occupancy is selective for promoters and actively transcribing transposons, rather than open chromatin in general.

**Fig. 4.**
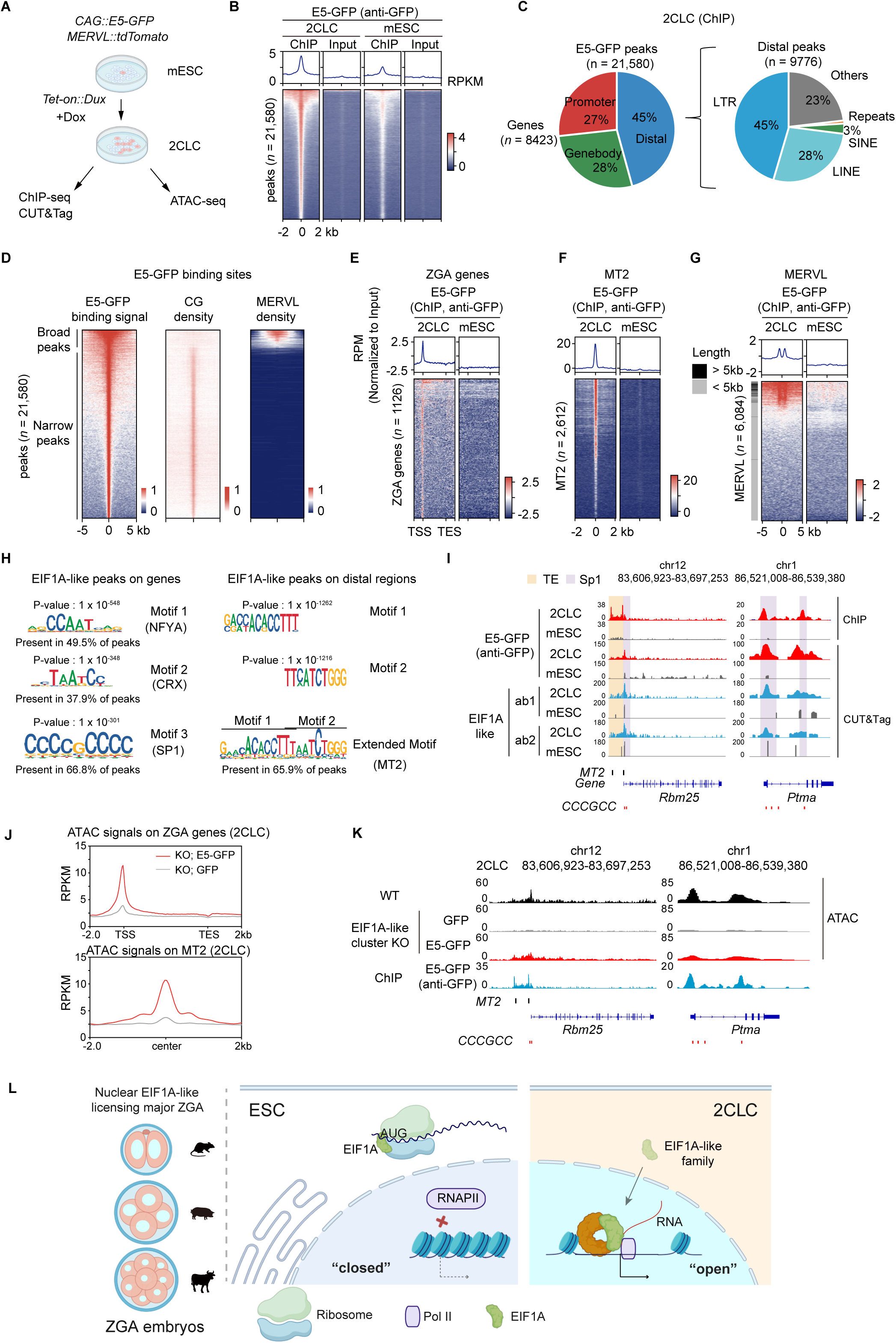
EIF1A-like proteins bind to chromatin associated with ZGA genes. (**A**) Schematic of ChIP–seq, CUT&Tag and ATAC–seq in mESCs and Dux-induced 2CLCs expressing E5-GFP. (**B**) ChIP–seq signal at E5-GFP peaks (*n* = 21,580) in 2CLCs and mESCs. (**C**) Genomic distribution of E5-GFP peaks (left) and retrotransposon composition of distal peaks (right). (**D**) E5-GFP peaks ranked by signal, shown with CG density and MERVL density. (**E** to **G**), E5-GFP ChIP–seq signal at ZGA genes (E), MT2 (F) and MERVL (G) elements in 2CLCs and mESCs. (**H**) Motifs enriched at EIF1A-like peaks on genes (left) and distal regions (right), with the closest matching transcription factors and prevalence among peaks. (**I**) ChIP–seq and CUT&Tag tracks at two representative loci (Rbm25, Ptma). (**J**) ATAC–seq signal at ZGA genes and MT2 elements in cluster-KO 2CLCs rescued with GFP or E5-GFP. (**K**) ATAC–seq and E5-GFP ChIP–seq tracks at the loci in i, in wild-type and rescued cluster-KO cells. (**L**) Model: nuclear EIF1A-like proteins license major ZGA.

CUT&Tag with anti-GFP antibody and two antibodies against endogenous EIF1A-like proteins independently reproduced the ChIP-seq pattern (fig. S7, H and I). The three datasets yielded 15,478 shared peaks, extensively overlapped ChIP-seq peaks and mapped to similar regions (fig. S7, J to I). EIF1A-like occupancy was especially strong at promoters of genes activated in 2-cell embryos, including housekeeping genes (Fig. 4E and fig. S8, A and B). These signals appeared in 2CLCs but not in mESCs or IgG controls and were confirmed by CUT&Tag pattern. Thus, EIF1A-like proteins occupy the promoters of genes activated during 2C-like induction.

EIF1A-like proteins also occupied ZGA-associated retrotransposons. Activation of MERVLs and MT2 is tightly associated with ZGA in 2C embryos and DUX-induced 2CLCs(*9, 29*). Nearly all MT2 sites were enriched for EIF1A-like signal, and approximately 20% of MERVLs were occupied, with a clear bias towards full-length or young elements (Fig. 4, F and G and fig. S8, C and D). Strikingly, promoter-proximal peaks were enriched for NFYA1, CRX, and SP1 motifs, whereas distal peaks contained an extended MT2-like motif(*12, 51, 52*) (Fig. 4, H and I and fig. S8E). At *ZSCAN4* and *REFEE* loci, occupancy concentrated at promoters that are frequently MT2-derived(*53, 54*) (fig. S8. F and G); at *Rbm25* and *Ptma*, it coincided with GC-rich SP1 motifs(*16, 52, 55, 56*) (Fig. 4H).

To test whether this pattern occurs in 2-cell embryos, we performed Stacc-seq with an antibody against endogenous EIF1A-like proteins and identified 29,359 peaks(*57*). Their genomic distribution closely matched that in 2CLCs, and 60% of occupied genes were also bound by RNA polymerase II (fig. S9, A and B). EIF1A-like proteins and polymerase II showed similar profiles across shared loci in embryos and 2CLCs, whereas no comparable signal was detected in mESCs or at embryonic enhancers (fig. S9, C and D). To assess direct DNA binding, we performed EMSA with recombinant EIF1A-like proteins (fig.S9E). Purified EIF1AD8 bound dsDNA efficiently, and increasing protein concentrations induced a super shift, consistent with cooperative binding. These results support a model wherein EIF1A-like proteins regulate ZGA genes through direct chromatin interaction.

### EIF1A-like proteins maintain local chromatin accessibility of ZGA genes

To investigate how promoter-bound EIF1A-like proteins regulate transcription, we performed ATAC-seq on tdTomato-positive cells sorted from *Eif1a-*like cluster-deletion mESCs expressing either GFP or E5-GFP upon *Dux* induction (fig. S10A). ATAC-seq showed that accessibility at ZGA-gene promoters and MT2 elements was markedly reduced in GFP-expressing knockout cells but restored by E5-GFP (Fig. 4J). The same relationship was evident at *Rbm25, Ptma, Zscan4,* and *REFEE* loci (Fig. 4K and fig. S10, B and C). Notably, accessibility changes aligned precisely with sites of EIF1A-like occupancy (fig. S10D), suggesting that EIF1A-like binding may shape local chromatin opening. Depletion in embryos similarly reduced accessibility at ZGA promoters and MT2 loci without altering global accessibility (fig. S10E). EIF1A-like proteins therefore act locally to establish or maintain accessible chromatin at their target promoters and retrotransposons.

## DISCUSSION

By placing divergent genomes in a common cytoplasm, interspecific hybrid embryos revealed a layer of mammalian ZGA that conventional species comparisons cannot resolve. Foreign oocytes activated sperm genomes separated by more than 60 million years of evolution, and hybrids contain a mouse genome adopted mouse-like ZGA timing. This cross-species framework uncovered a core set of genes reproducibly activated across diverse embryonic contexts, with the *Eif1a-like* family among the most prominent. Loss-of-function experiments in multiple species establish EIF1A-like proteins as conserved requirements for ZGA and early development across mammals. These results argue against the commonly held view that ZGA regulation is species-specific.

EIF1A-like proteins are conventionally associated with translation initiation, yet their critical activity during ZGA is nuclear. Their nuclear translocation tracks major ZGA in mouse, pig and cattle. Consistently, this pattern is reproduced in 2CLCs but not mESCs. Functionally, forced nuclear localization partially restored early embryonic development, whereas nuclear exclusion does not. Once in the nucleus, EIF1A-like proteins occupy ZGA promoters and ZGA-associated retrotransposons, where they are required for accessibility at these sites. Therefore, EIF1A-like proteins meet the operational definition of a ZGA licensing factor: creating a permissive chromatin state without being sufficient to initiate the program on its own^34,53^ (Fig. 4L). This unexpected nuclear function of a canonical translation factor unveils a previously unappreciated paradigm of transcriptional regulation governing the maternal-to-zygotic transition. This suggests that mammals may share a conserved strategy of producing these licensing factors *de novo* at the onset of ZGA.

How EIF1A-like proteins open chromatin remains unresolved. Their occupancy is sharply focused on promoters and transcribed retrotransposons, and their depletion reduces accessibility at these loci. Consistently, EIF1A associated motifs are highly GC-rich, matching known features of ZGA gene promoters. They are also enriched for several pioneer transcription factors (TFs) known important for ZGA. Interestingly, the OB domain of EIF1A is frequently found in other DNA binding proteins, echoing our EMSA results *in vitro*. The direct DNA binding fits well with the fact that EIF1A-like proteins maintain an invariant NLS, conserved from mammals to nematodes, raises the possibility that the nuclear role is not a recent but rather an ancient function, perhaps as old as translation. One possibility is that *de novo* synthesis of EIF1A-like proteins couples the first productive zygotic transcripts to a second, broad wave of genome activation. In mouse, expansion of *Eif1a-like* locus may provide the dosage needed to act as early as 2-cell to activate thousands of promoters. Such a feed-forward circuit would allow the awakening genome to reinforce its own transcriptional competence and make the transition away from maternal control increasingly irreversible(*6*). This model is attractive but provisional, requiring further tests of protein synthesis, nuclear import and promoter opening in the same embryos (Fig. 4L).

Together, our results separate the identity of a conserved licensing machinery from the species-specific timing of its deployment. This distinction explains how mammalian ZGA can evolve rapidly in sequence and schedule while retaining a shared mechanistic core. Beyond ZGA, interspecies hybrid transcriptomics provides a general framework for dissecting cross-species regulatory compatibility in other developmental transitions. Key questions remain to be explored in the future: how EIF1A-like proteins cooperate with pioneer factors to activate transcription, how their nuclear and cytoplasmic activities are coordinated, what genomic feature encodes species-specific timing, and whether this licensing strategy extends beyond mammals.

## ACKNOWLEDGMENTS

We thank members of Zhao and Yu laboratory for discussion and comments during the study and preparation of the manuscript. We thank the Animal Center at Institute of Biophysics, Chinese Academy of Sciences, Institute of Zoology, Chinese Academy of Sciences and the Laboratory Animal Welfare and Ethics Committee of Guangzhou Medical University for their support. We thank Jian Zhang for helpful discussions. We thank Yingchun Ouyang and Guosong Qing from Institute of Zoology for their help with ICSI. We thank Bing Zhu from Institute of Biophysics for sharing the inducible 2CLC reporter line.

## Author contributions

Conceptualization, supervision and writing – original draft, Y.Y., D.Q., K.C., N.Y., W.X., J.G.Z.; methodology, D.Q., D.F.B., Z.H.Q., M.W. and Y.Y.; investigation, D.Q., Z.H.Y., D.F.B., Z.C.Z., W.W.L., Z.L.W., Y.L.L., L.J.D., Y.N.D., J.M.C., Z.H.C., J.C.Z., S.Q.G., J.C.D., Y.K.J., X.Z.Z., T.Y.Z., Y.B.W., Y.H.Y., M.H., Q.Y.Z. and S.W.; formal analysis, Z.H.Q., D.Q., X.Y.W., X.H.F., Z.Y.Z., and Z.Y.R.; resources, J.G.Z., W.X., W.L., C.L., L.S., W.H.Y., Q.R.K., S.R.G., Y.K.D., S.W., J.L., Y.Q., Y.F.W., M.Q.D. and Y.Y.; writing – review and editing, D.Q., Z.H.Q., X.H.L., D.F.B., Y.N.D., W.W.L., Q.Q.S., T.J.Z. and Y.Y.; funding acquisition, Y.N., K.C., W.X., J.G.Z. and Y.Y.

## Data and materials availability

The sequencing data have been deposited to the Gene Expression Omnibus (GEO) with accession number GSE328869, GSE328840 and GSE328839. The mass spectrometry proteomics data have been deposited to the ProteomeXchange Consortium via the PRIDE partner repository with the dataset identifier PXD078112.

## Declaration of generative AI and AI-assisted technologies in the writing process

During the preparation of this work, the authors used ChatGPT and Kimi to improve the English. After using these tools, the authors reviewed and edited the content as needed and took full responsibility for the content of the publication.

---

**Figure S1.**
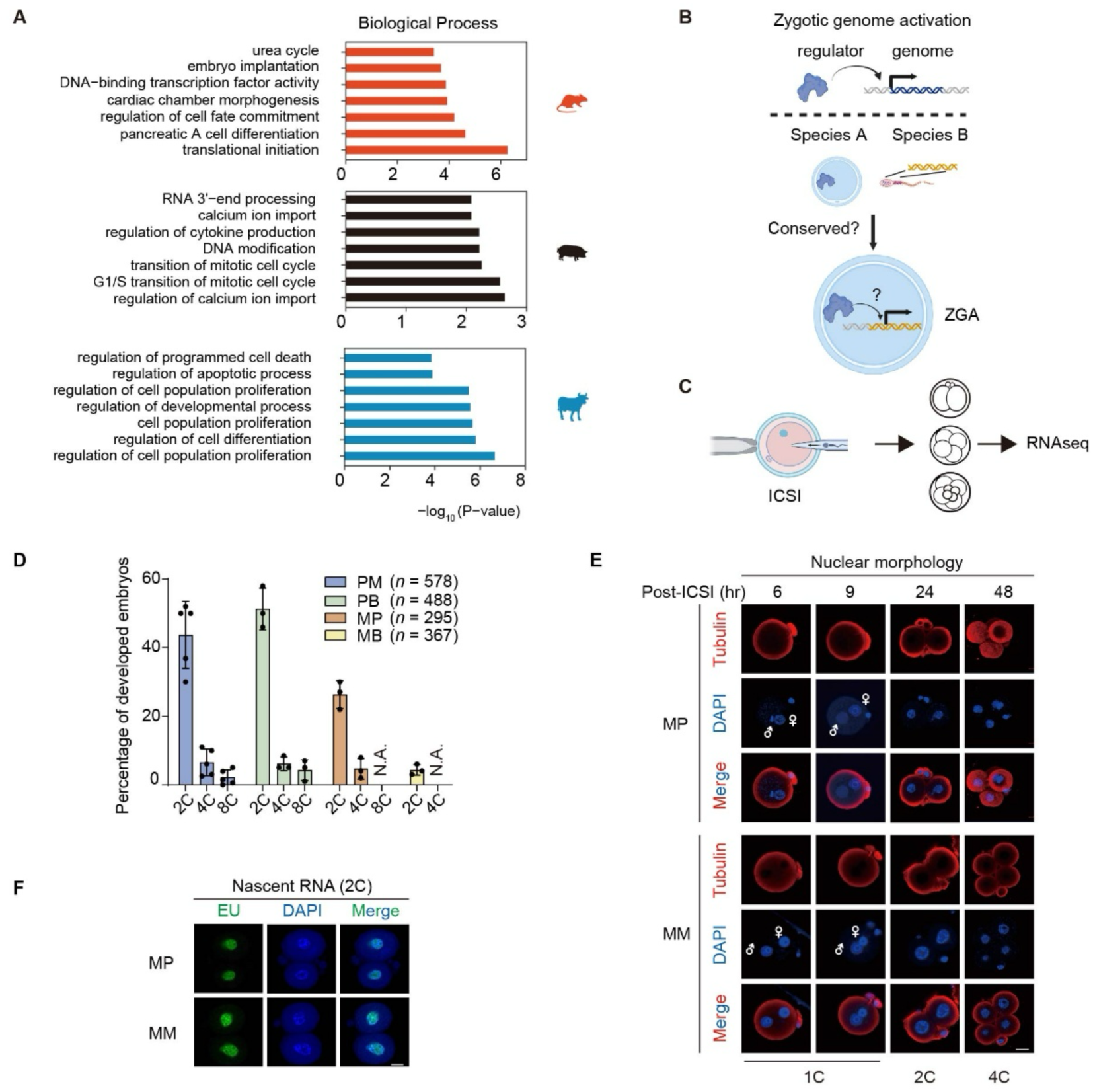
An interspecies hybrid assay for testing conservation of ZGA regulation. (**A**) Gene Ontology enrichment of ZGA genes in mouse, pig and cattle embryos. (**B**) Schematic of the cross-species assay: the genome of species A is placed under the maternal machinery of a species B oocyte to test whether ZGA regulation is conserved. (**C**) Experimental workflow. (**D**) Percentages of PM, PB, MP and MB embryos reaching the indicated cleavage stages. n, total embryo number. (**E**) Nuclear morphology of MP and MM embryos at the indicated hours post-ICSI, stained for tubulin and DNA (DAPI). Scale bar, 20 μm. (**F**) Nascent transcription at the two-cell stage in MP and MM embryos, detected by EU incorporation. Scale bar, 20 μm.

**Figure S2.**
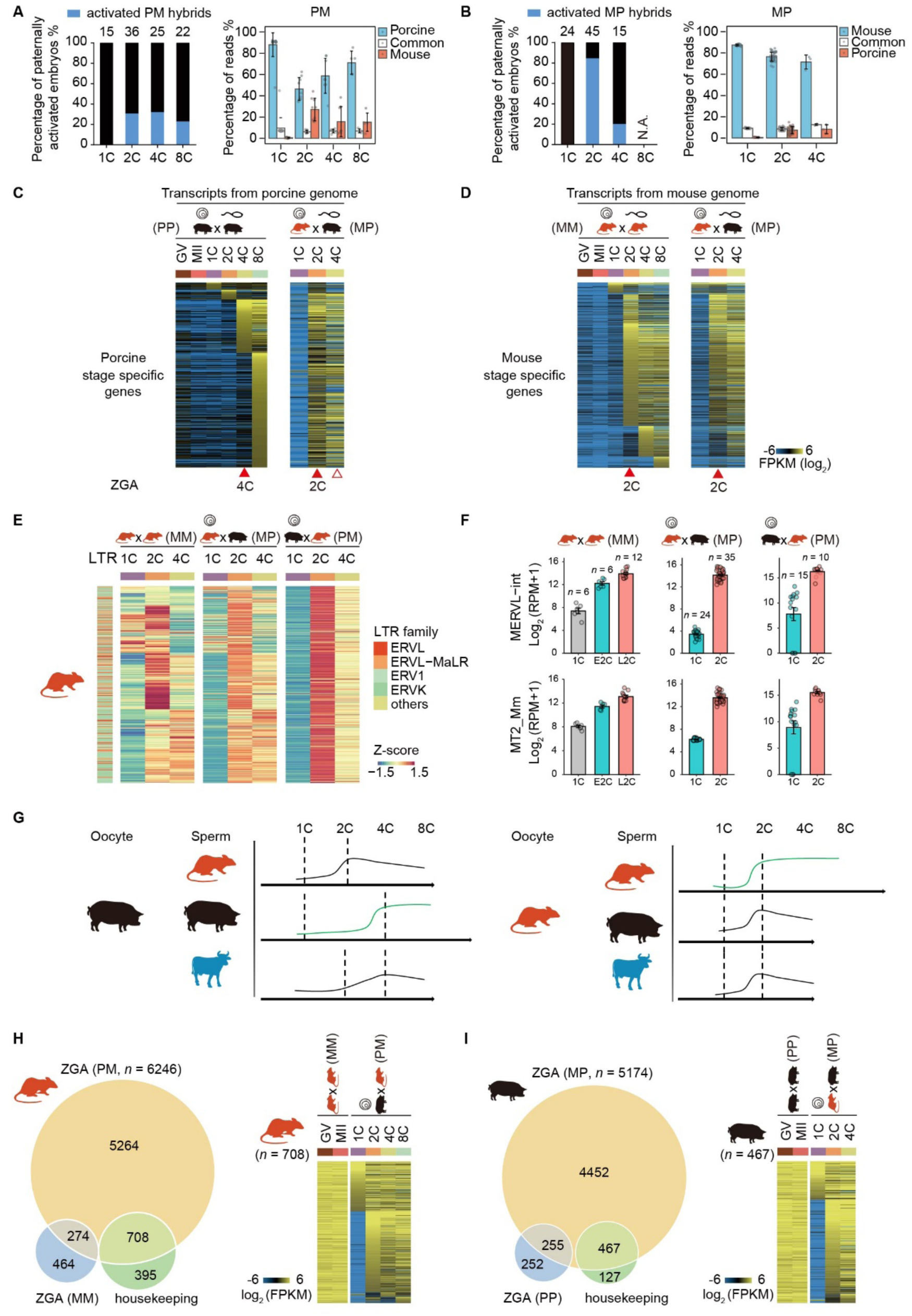
Faithful activation of the sperm genome in foreign oocytes. (**A** and **B**) Bar graph showing percentage of paternally activated hybrid embryos (Left) and reads mapping to either paternal, maternal, or both genomes (right) in various hybrid embryos: PM (A), MP (B) at different developmental stages, the number of embryos sequenced is shown above the bars. N.A., not available. (**C** and **D**) Heatmaps showing gene expression in MP hybrid embryos. Transcripts from the porcine genome (C) or the mouse genome (D) were compared to the corresponding stage-specific activated genes of the indicated normal embryos. (**E**) Expression of LTR retrotransposon families in MM, MP and PM embryos at the indicated stages (Z-score). (**F**) Expression of MERVL-int and MT2_Mm in MM, MP and PM embryos. n, embryo number. (**G**) Schematic of hybrid embryo construction by ICSI and the onset of paternal genome activation in each combination. (**H** and **I**) Overlap of ZGA genes activated in PM (H) and MP (I) hybrids with ZGA and housekeeping genes of the corresponding natural embryos (MM, PP); heatmaps show expression of the overlapping genes across stages.

**Figure S3.**
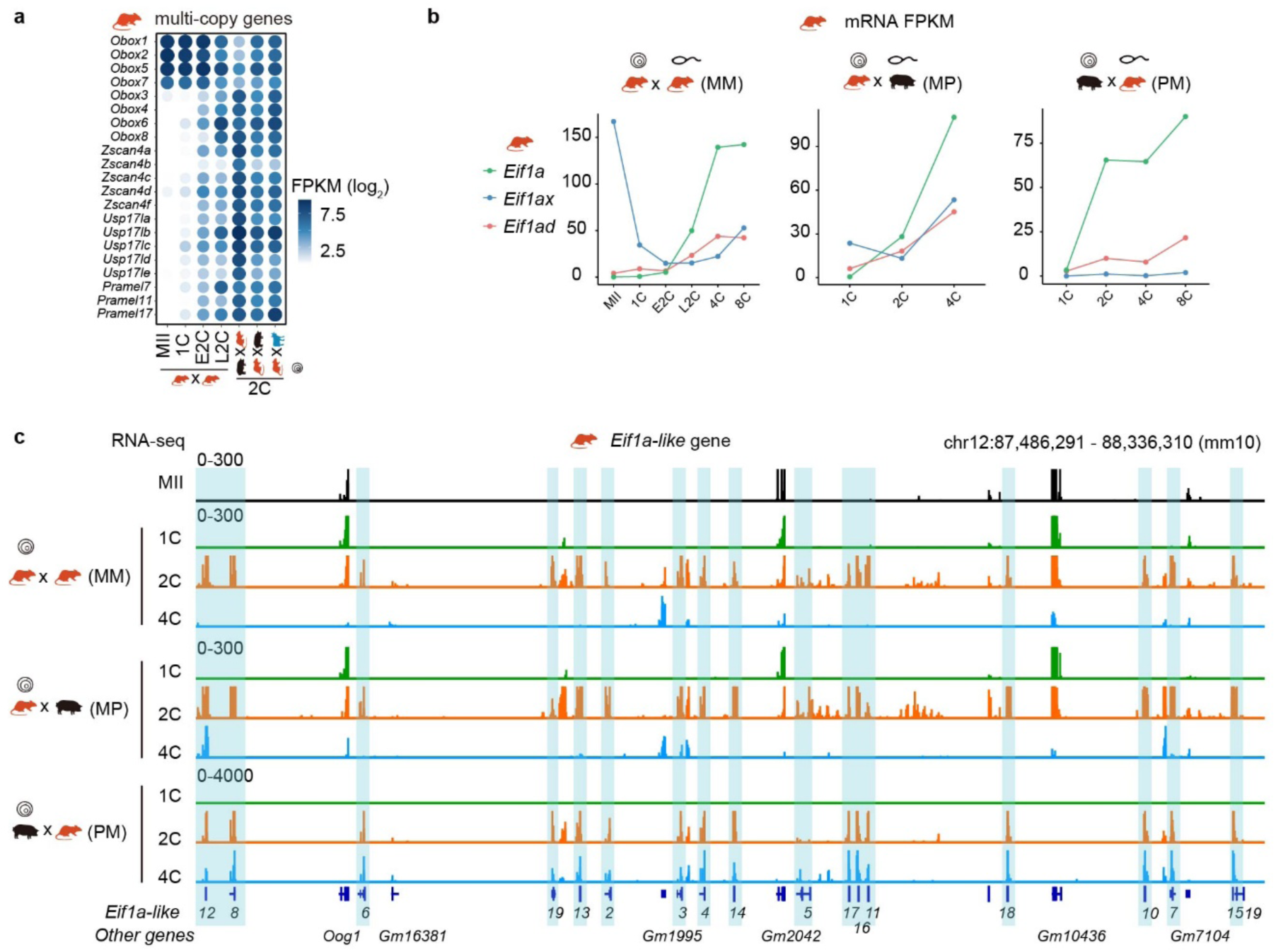
Zygotic activation of *Eif1a*-family genes in natural and hybrid embryos. (**A**) Expression of multi-copy ZGA genes in mouse MII oocytes, cleavage-stage embryos and two-cell hybrid embryos. (**B**) mRNA levels of *Eif1a*, *Eif1ax* and *Eif1ad* in MM, MP and PM embryos at the indicated stages. (**C**) RNA-seq tracks of the *Eif1a-like* gene cluster (chr12, mm10) in MII oocytes and MM, MP and PM embryos. Numbered *Eif1a-like* genes are highlighted.

**Figure S4.**
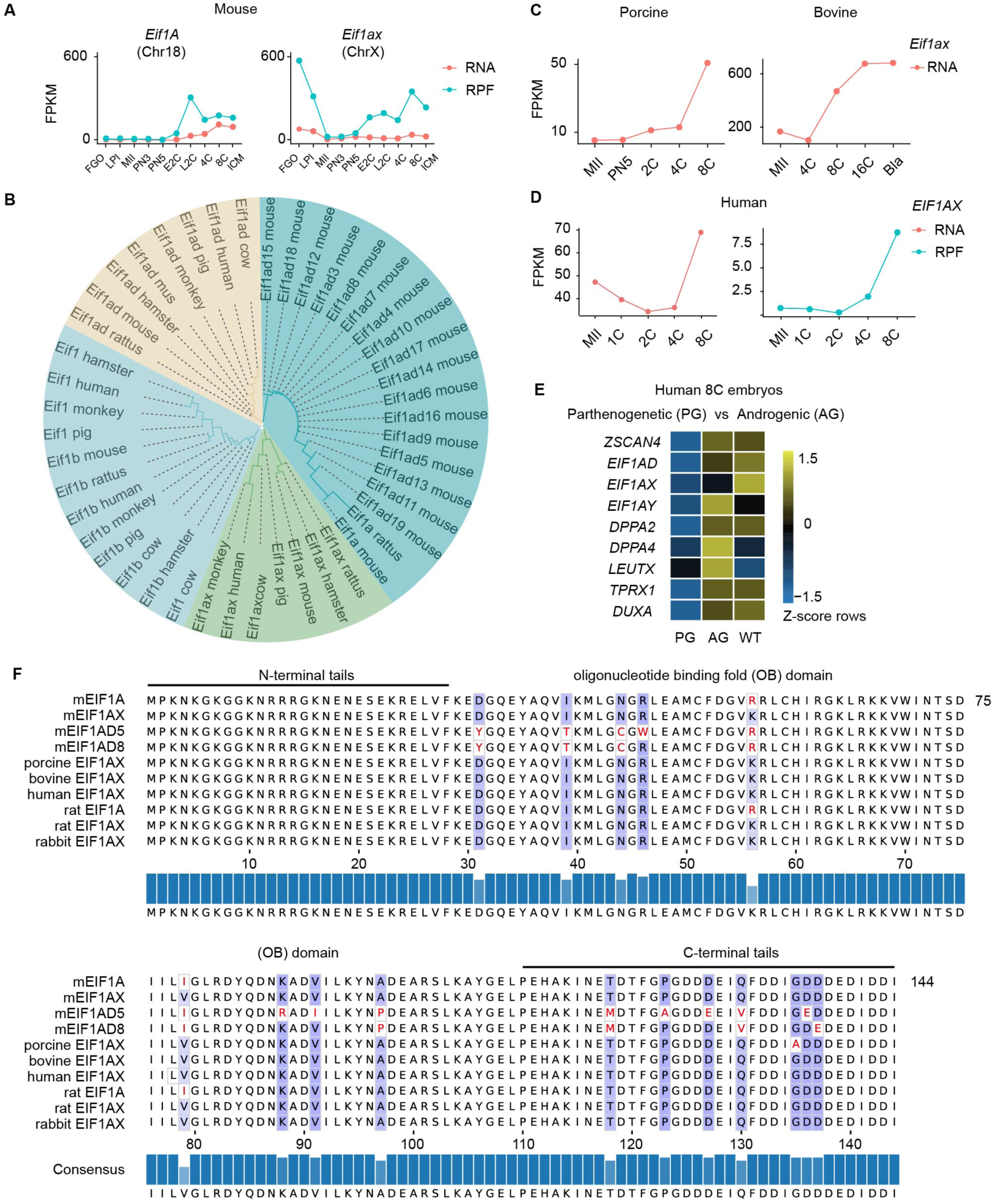
Evolution and expression of the EIF1A family across mammals. (**A**) RNA-seq (red) and ribosome protect fragment (RPF; cyan) expression dynamics of canonical *Eif1a* (Chr18) and *Eif1ax* (ChrX) across mouse oocyte and preimplantation stages. (**B**) Phylogenetic tree of EIF1A-family proteins across mammalian species. Copy-number expansion of Eif1ad is specific to mouse. (**C**) Eif1ax mRNA levels in porcine and bovine oocytes and embryos at the indicated stages. (**D**) EIF1AX mRNA (RNA) and ribosome-protected fragment (RPF) levels in human oocytes and embryos. (**E**) Expression of ZGA genes in human eight-cell parthenogenetic (PG), androgenetic (AG) and wild-type (WT) embryos (row Z-score). (**F**) Amino acid sequence alignment of EIF1A-family proteins. The N-terminal tail, oligonucleotide-binding (OB) domain and C-terminal tail are indicated; consensus is shown below.

**Figure S5.**
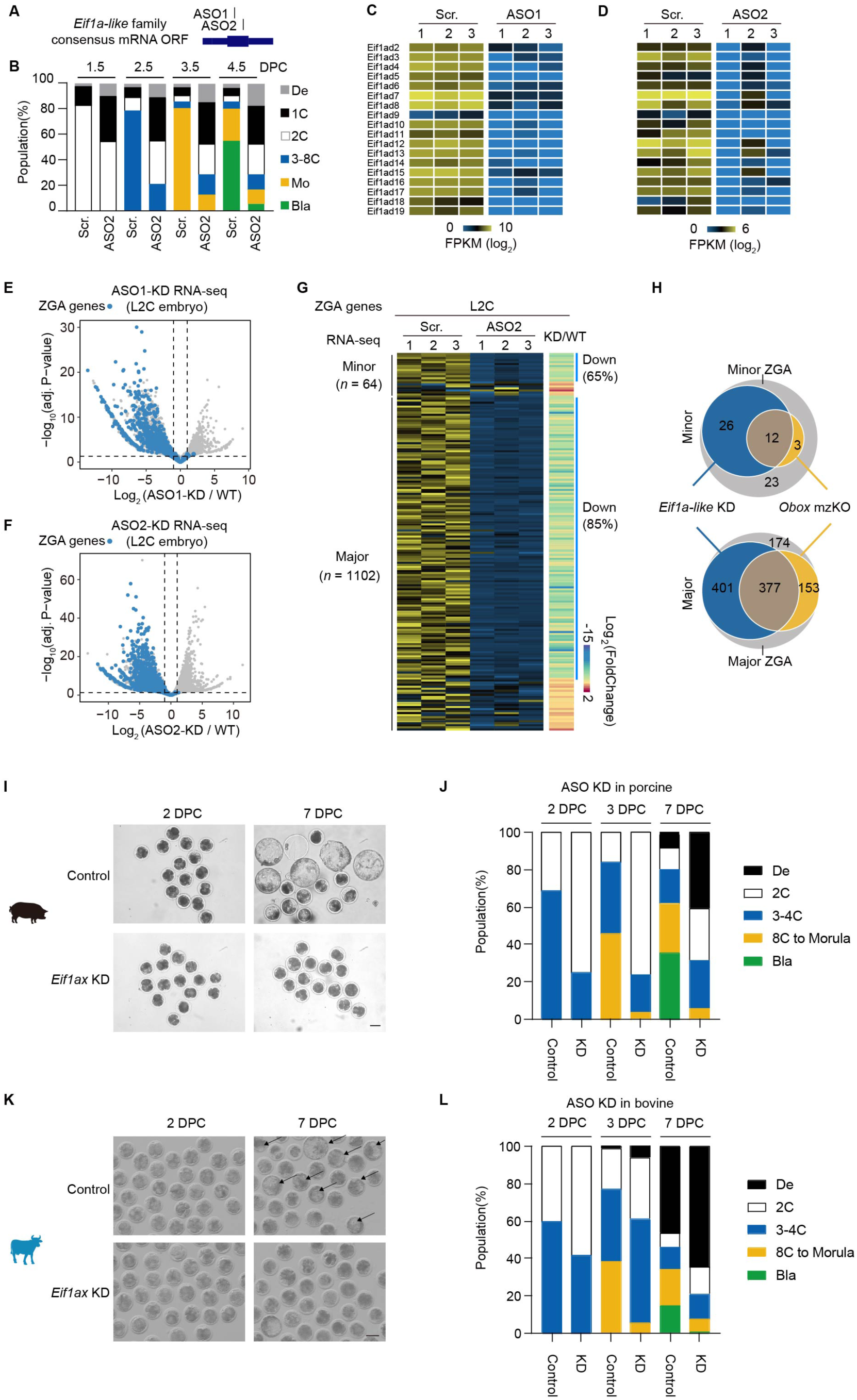
ASO-mediated depletion of *Eif1a* family transcripts impairs ZGA and development. (**A**) Schematic of antisense oligonucleotides (ASO1, ASO2) targeting the consensus open reading frame of *Eif1a*-like family mRNAs. (**B**) Developmental stage distribution of scrambled-control (Scr.) and ASO2-injected embryos at the indicated days post coitum. (**C** and **D**) Expression of *Eif1a-like* family members in Scr.- and ASO1 (C) or ASO2-injected (D) late two-cell embryos. (**E** and **F**) Volcano plots of RNA-seq in ASO1 (E) and ASO2-KD (F) late two-cell embryos; ZGA genes are highlighted in blue. (**G**) Heatmap of RNA-seq expression for minor ZGA and major genes in L2C embryos treated with Scr. and ASO2. (**H**) Overlap of genes downregulated in *Eif1a-like* KD with minor and major ZGA genes and with *Obox* mzKO targets. (**I** to **L**), Representative bright-field images of control and *Eif1ax*-depleted porcine (I) and bovine (K) embryos at 2 and 7 DPC. Scale bars, 100 μm. Developmental stage distribution of control and ASO-injected porcine (J) and bovine (L) embryos at 2, 3 and 7 DPC.

**Figure S6.**
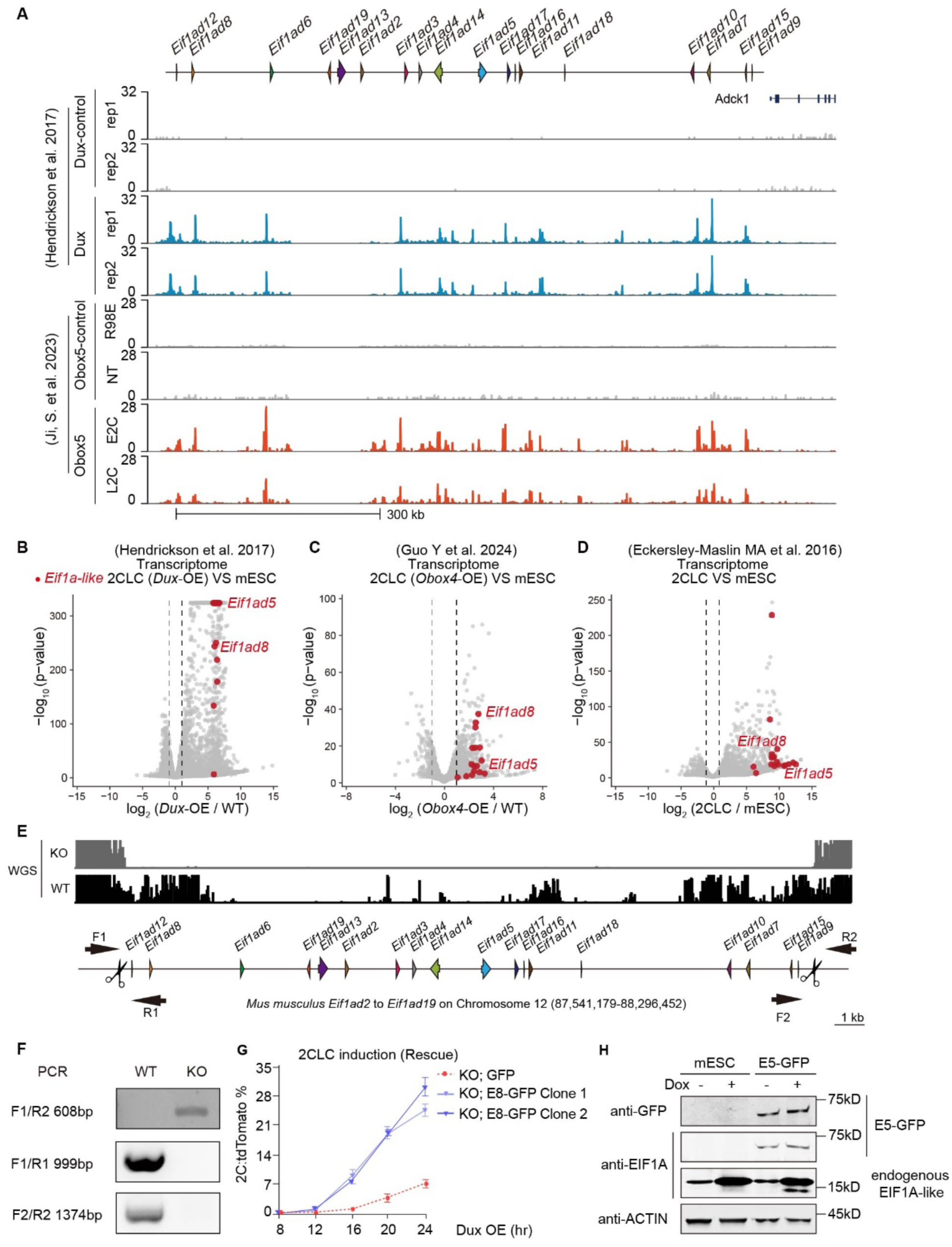
The *Eif1a-like* cluster is required for 2CLC reprogramming. (**A**) DUX and OBOX5 occupancy at the mouse *Eif1a-like* gene cluster. (**B** to **D**) Reanalysis of published transcriptomes comparing 2CLCs with mESCs: Dux overexpression (B), Obox4 overexpression (C) and spontaneous 2CLCs (D). Eif1a-like genes are highlighted in red. (**E**) Whole-genome sequencing (WGS) of wild-type (WT) and cluster-knockout (KO) mESCs at the Eif1ad2–Eif1ad19 locus. Arrows, genotyping primers. (**F**) PCR genotyping of WT and KO mESCs. (**G**) 2CLC induction after *Dux* overexpression in KO mESCs expressing GFP or E8-GFP (two clones). (**H**) Western blot validation of E5-GFP expression (anti-GFP, ∼72 kDa) and endogenous EIF1A-LIKE (anti-EIF1A, ∼17 kDa) in mESCs and E5-GFP cells ± Dox. ACTIN serves as loading control.

**Figure S7.**
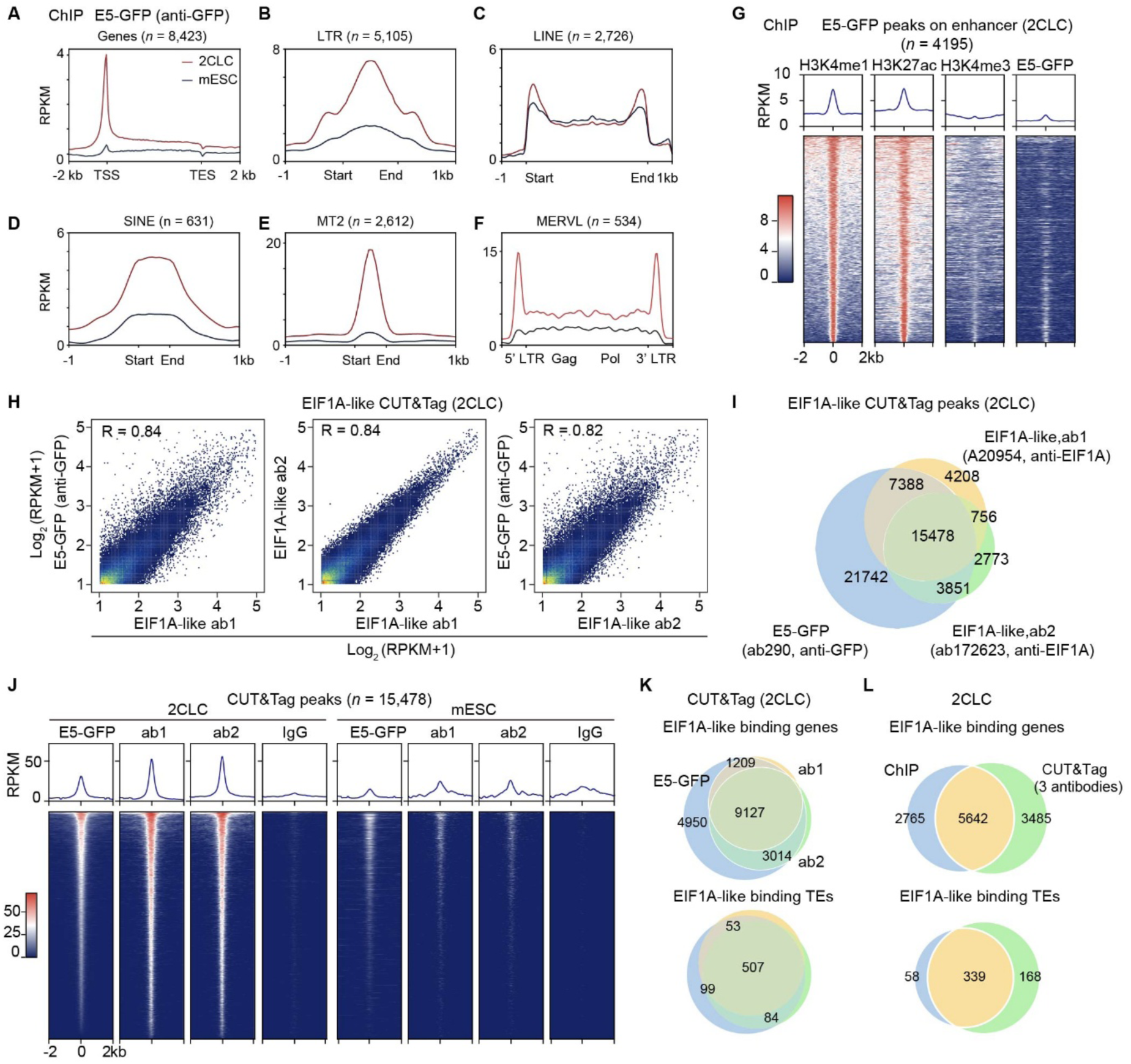
Genome-wide EIF1A-like occupancy mapped by ChIP–seq and CUT&Tag. (**A** to **F**) ChIP–seq metagene profiles of E5-GFP (anti-GFP) in two-cell-like cells (2CLCs) and mouse embryonic stem cells (mESCs) over genes (A), LTR (B), LINE (C) and SINE (D) retrotransposons, MT2 (E) and MERVL (F) elements. n, detected element number. (**G**) Metagene profiles and heatmaps of H3K4me1, H3K27ac, H3K4me3 and E5-GFP ChIP-seq signals centered on 2CLC enhancer regions. (**H**) Pairwise correlations of CUT&Tag signal between E5-GFP (anti-GFP) and two anti-EIF1A antibodies (ab1, ab2) in two-cell-like cells (2CLCs). R, Pearson correlation coefficient. (**I**) Overlap of CUT&Tag peaks called with each antibody in 2CLCs. (**J**) Heatmaps and metagene profiles of CUT&Tag signal at the 15,478 shared peaks in 2CLCs and mouse embryonic stem cells (mESCs). IgG, negative control. (**K**) Overlap of EIF1A-like-bound genes and transposable elements (TEs) across the three antibodies. (**L**) Overlap of EIF1A-like-bound genes and TEs between ChIP–seq (E5-GFP) and CUT&Tag (three antibodies combined).

**Figure S8.**
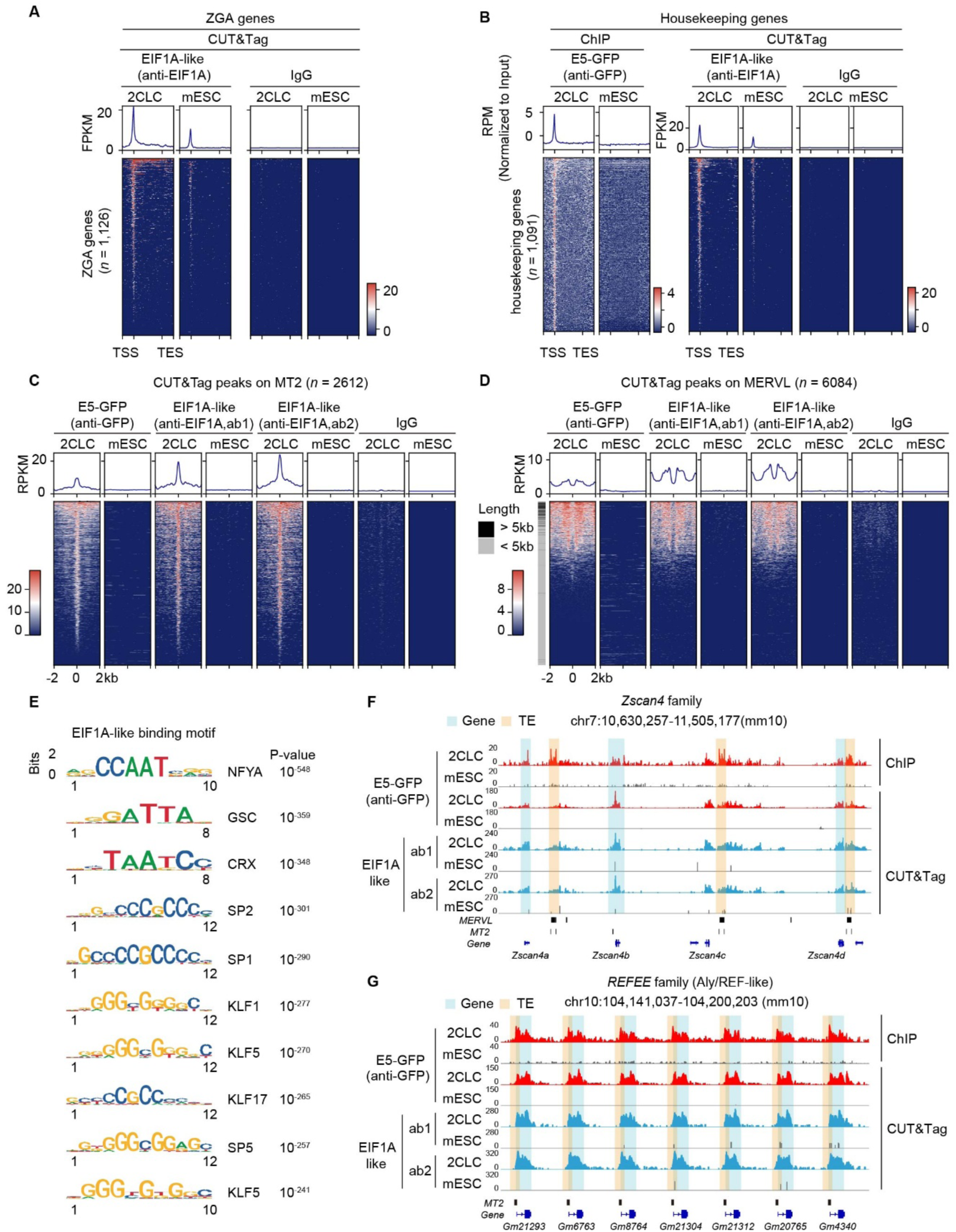
Specificity of EIF1A-family occupancy at ZGA-gene promoters. (**A** and **B**) CUT&Tag or ChIP–seq heatmaps and metagene profiles at ZGA genes (A) and housekeeping genes (B) in 2CLCs and mESCs. (**C** and **D**) CUT&Tag signal at MT2 (C) and MERVL (D) elements in 2CLCs and mESCs. In d, elements are color-coded by length. (**E**) Top motifs enriched at EIF1A-like-bound loci. (**F** and **G**) ChIP–seq and CUT&Tag tracks at the *Zscan4* family locus (F) and the *REFEE* family locus (G). Genes and transposable elements (TEs) are indicated below.

**Figure S9.**
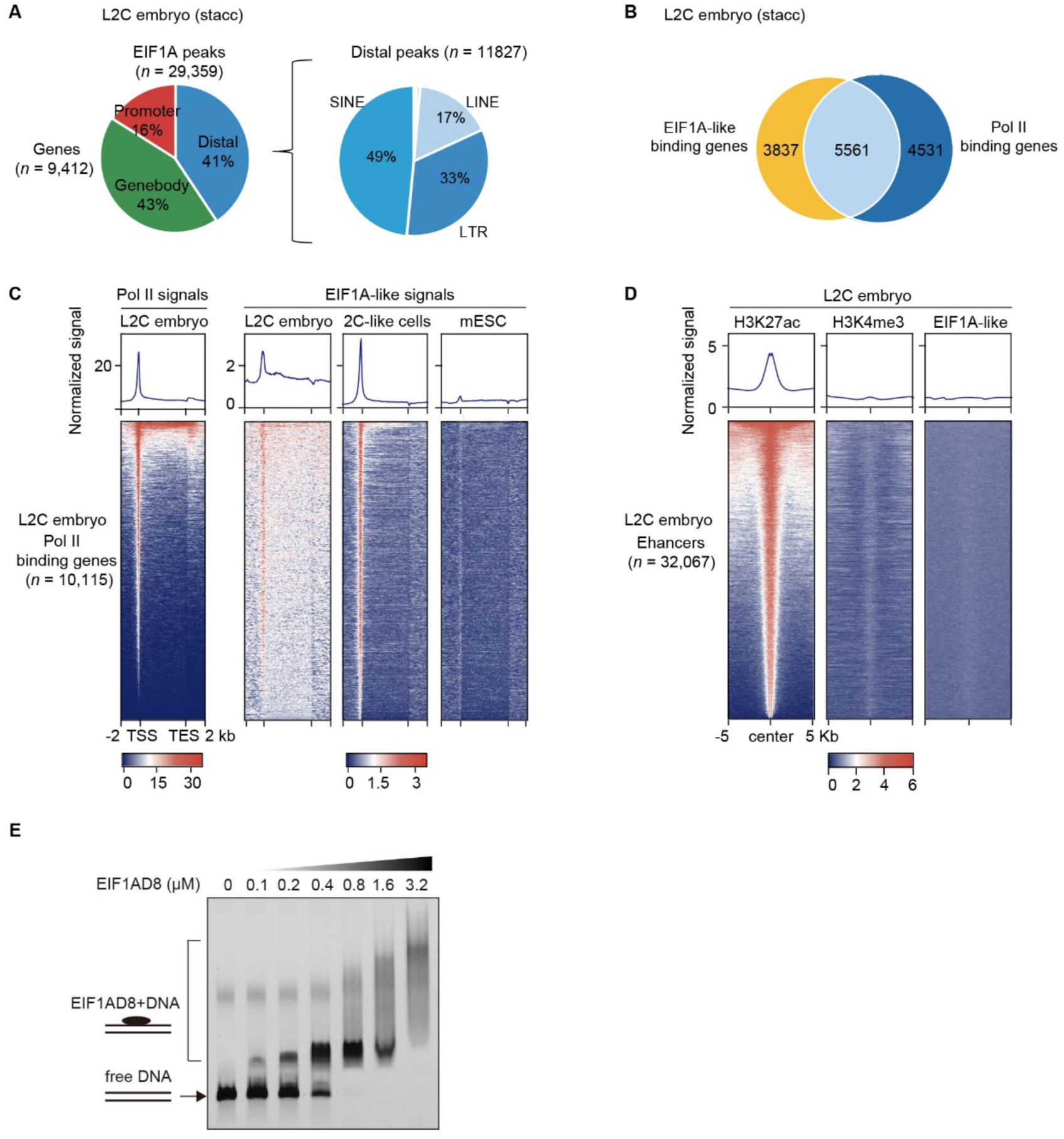
Stacc-seq confirms EIF1A-like occupancy at ZGA loci in 2-cell embryos. (**A**) Genomic distribution of EIF1A-like Stacc-seq peaks in late two-cell (L2C) embryos (left) and retrotransposon composition of distal peaks (right). (**B**) Overlap of EIF1A-like-bound genes and RNA polymerase II (Pol II)-bound genes in L2C embryos. (**C**) Pol II and EIF1A-like signals at Pol II-bound genes in L2C embryos, with EIF1A-like signals in 2C-like cells and mESCs for comparison. (**D**) H3K27ac, H3K4me3 and EIF1A-like signals at L2C embryo enhancers. (**E**) Electrophoretic mobility shift assay (EMSA) showing dose-dependent binding of recombinant EIF1AD8 to DNA. Increasing concentrations of purified EIF1AD8 (0, 0.1, 0.2, 0.4, 0.8, 1.6 and 3.2 μM) were incubated with 50 nM DNA probe.

**Figure S10.**
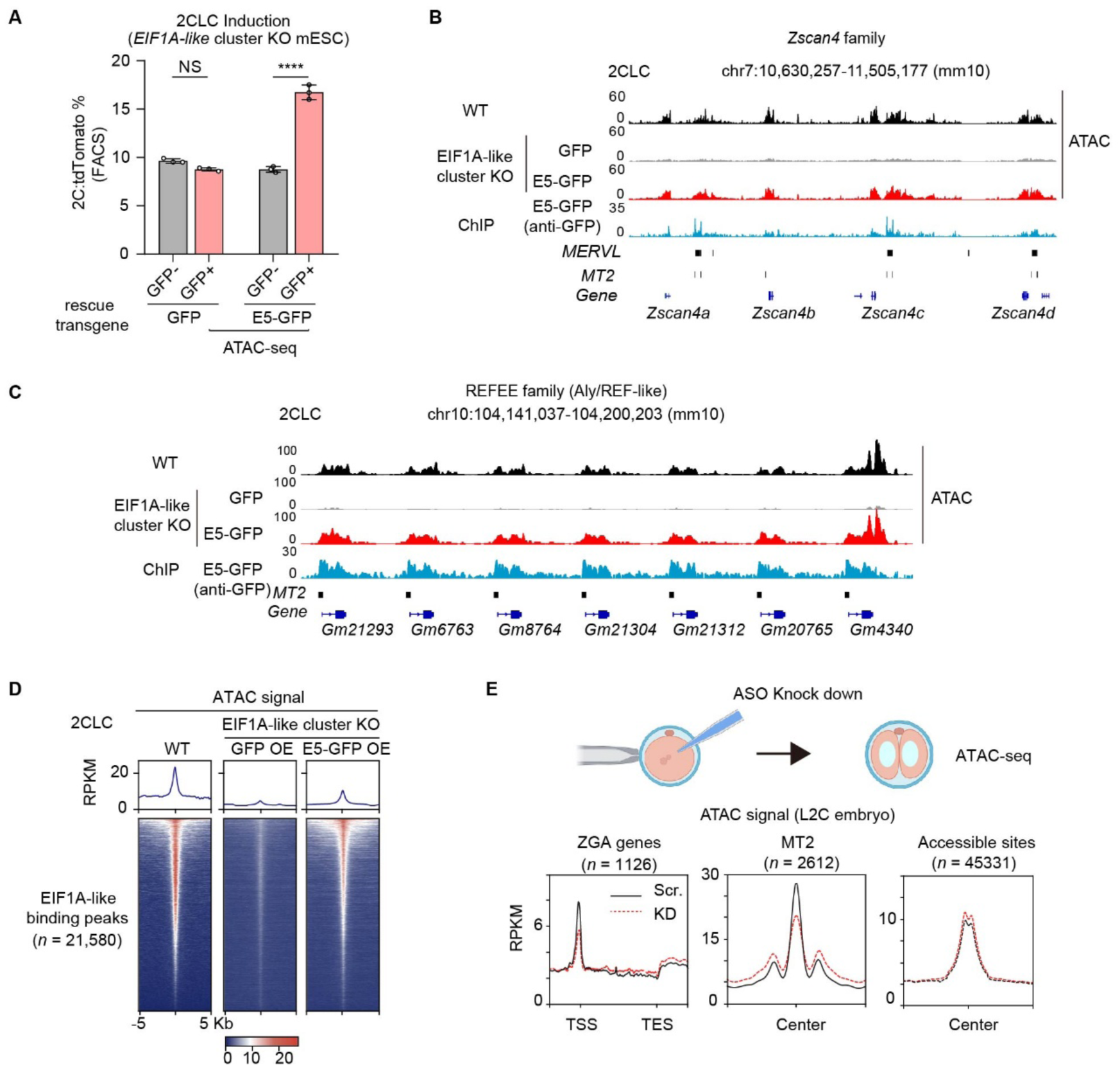
EIF1A family proteins are required for local chromatin accessibility. (**A**) 2CLC induction in Eif1a-like cluster-KO mESCs rescued with GFP or E5-GFP (FACS). NS, not significant; ****P < 0.0001. GFP-positive and E5-GFP-positive 2CLCs were sorted and used for ATAC-seq in e–g. (**B** and **C**) ATAC-seq and E5-GFP ChIP–seq tracks at the Zscan4 family (B) and REFEE-family (C) loci in wild-type (WT) and cluster-KO cells rescued with GFP or E5-GFP. (**D**) ATAC-seq signal at EIF1A-like-bound peaks in the indicated samples. (**E**) ATAC-seq signal at ZGA genes, MT2 elements and all accessible sites in control (Scr.) and ASO-depleted (KD) late two-cell embryos.

## Methods

### Ethics statement

The study protocols were approved by the Biosafety, Ethics and Laboratory Animal Management Committees of the Institute of Biophysics, Chinese Academy of Sciences (approval no. SYXK2020057), the Institute of Zoology, Chinese Academy of Sciences (approval no. IOZ20180075) and the Laboratory Animal Welfare and Ethics Committee of Guangzhou Medical University (approval no. GY2024-558). All efforts were made to minimize animal suffering and to reduce the number of animals used.

### Fertilization and embryo collection

Female B6D2F1 or C57BL/6 mice aged 3-4 weeks were first injected with 5 IU pregnant mare serum gonadotropin (PMSG), followed 48 hours later by a second injection of 5 IU human chorionic gonadotropin (hCG). Cumulus-oocyte complexes were collected from the oviducts 13-14 hours after hCG injection. Fresh sperm from B6D2F1 or C57BL/6 males were capacitated by incubation in human tubal fluid (HTF) medium at 37°C for at least 1 hour. In vitro fertilization (IVF) was performed by incubating cumulus-oocyte complexes with capacitated sperm for 4-6 hours at 37°C in an atmosphere of 5% CO2. Fertilized embryos were washed once in HTF medium and then transferred to KSOM medium for further culture until collection. Embryos were collected at the following time points after fertilization: zygotes at 12 hours, early 2-cell (E2C) embryos at 20 hours, mid 2-cell (M2C) embryos at 28 hours, late 2-cell (L2C) embryos at 34 hours, 4-cell (4C) embryos at 48 hours, 8-cell (8C) embryos at 52 hours and blastocysts at 96 hours.

### In vitro maturation of porcine oocytes

Porcine ovaries were collected from prepubertal gilts at a local abattoir. These gilts were the progeny of a crossbred lineage involving Duroc boars and gilts descended from Landrace sires and Yorkshire dams. The ovaries were transported to the laboratory in saline at 37°C. Follicles measuring 3-6 mm in diameter were aspirated using an 18-gauge needle attached to a 10 ml syringe. Cumulus-oocyte complexes (COCs) in the follicular fluid were allowed to settle by gravity at 37°C. The COCs were then washed three times in HEPES-buffered Tyrode’s medium (6.663 g NaCl, 0.237 g KCl, 0.168 g NaHCO_3_, 0.041 g NaH_2_PO_4_, 1.868 ml sodium lactate, 0.102 g MgCl_2_·6H_2_O, 2.383 g HEPES, 0.065 g penicillin G, 0.010 g phenol red, 0.294 g CaCl_2_·2H_2_O, 2.186 g sorbitol, 0.025 g gentamicin, 0.022 g sodium pyruvate, and 1,000 ml Milli-Q H_2_O) containing 0.01% PVA at 37°C. Only COCs with multiple layers of intact cumulus cells and homogeneous ooplasm were selected for in vitro maturation (IVM). After three washes in IVM medium (TCM 199, Gibco BRL, supplemented with 0.1% PVA, 3.05 mM D-glucose, 0.91 mM sodium pyruvate, 1 μg/ml gentamicin, 0.57 mM cysteine, 0.5 μg/ml luteinizing hormone, 0.5 μg/ml follicle-stimulating hormone, and 10 ng/ml epidermal growth factor), groups of 70-80 COCs were transferred to wells of four-well culture plates (Nunc), each containing 500 μl IVM medium overlaid with 350 μl mineral oil. The COCs were cultured for 42-44 hours at 38.5°C in a humidified atmosphere of 5% CO_2_ in air. The mature COCs were then vortexed for 4 minutes in 0.1% hyaluronidase in HEPES-buffered Tyrode’s medium supplemented with 0.01% PVA to disperse the cumulus cells. Only oocytes that had reached maturation, as indicated by extrusion of the first polar body and uniform cytoplasm, were selected for further development. For IVF, porcine MII oocytes were washed three times in modified Tris-buffered fertilization medium (10 mM Tris containing 2 mg/ml BSA and 2 mM caffeine). Approximately 30-35 oocytes were transferred into 50 μl droplets of fertilization medium covered with mineral oil and pre-equilibrated for 4 hours at 38.5°C in 5% CO_2_ in air. For intracytoplasmic sperm injection (ICSI), porcine MII oocytes were centrifuged at 12,000 rpm for 10 minutes in a 1.5 ml centrifuge tube containing 500 μl TL-HEPES medium. After centrifugation, oocytes with clear cytoplasm were selected for ICSI.

### Intracytoplasmic sperm injection

Intracytoplasmic sperm injection was performed using an inverted microscope (Nikon Eclipse Ti-S, Nikon, Tokyo, Japan) equipped with a 20x objective lens (Nikon LWD 20x) and micromanipulators (Eppendorf, Hamburg, Germany). A drop of M2 medium was placed at the center of a 60 mm dish and covered with oil. Each oocyte was oriented with the first polar body at the 6 or 12 o’clock position and injected from the 3 o’clock position. A spermatozoon was introduced into the oocyte cytoplasm using an injection pipette controlled by a piezo micromanipulator (PMAS-CT150, Prime Tech, Japan).

### Sperm preparation

Porcine semen straws from a single Duroc boar (Baijunda Corporation, China) were thawed in a 50°C water bath for 16 seconds. After thawing, the semen was washed three times by centrifugation in Dulbecco’s phosphate-buffered saline supplemented with 1 mg/ml BSA at 1900 g for 4 minutes. The spermatozoa were then resuspended in modified Tris-buffered fertilization medium to a concentration of 1 × 10^6^ cells/ml. For IVF, 50 μl of the sperm suspension was added to 50 μl fertilization droplets containing 30-35 oocytes, yielding a final sperm concentration of 0.5 × 10^6^ cells/ml. For ICSI, motile sperm were rendered immotile using an Ultrasonic Homogenizer (JY92-IIN) at power setting 5 for 3 minutes in an ice bath. The immotile sperm were then washed once by centrifugation through a mixture of 70% Percoll and 30% M2 medium at 3,000 g for 5 minutes, followed by one wash in M2 medium and resuspension in M2 medium.

Bovine semen straws from a single Chinese Holstein bull (Beijing Shou Nong Corporation, China) were thawed in a 37°C water bath for 30 seconds. Bovine sperm used for ICSI were processed as described above for porcine sperm. Fresh mouse semen was collected and processed similarly to porcine sperm for ICSI.

### Electrical activation of oocytes

Porcine oocytes at 2-4 hours after ICSI were activated with two direct-current pulses of 1.0 kV/cm for 20 microseconds each, separated by a 1-second interval, using a BTX Electro Cell Manipulator 200 in fusion medium containing 0.3 M mannitol, 1 mM CaCl_2_, 0.1 mM MgCl_2_, and 0.55 mM HEPES.

### Hybrid embryo culture and collection

Porcine oocytes fertilized with hybrid sperm by ICSI were transferred to four-well plates, each containing 500 μl PZM3 medium overlaid with 350 μl mineral oil. The plates were incubated at 38.5°C in a humidified atmosphere containing 5% CO_2_. After electrical activation, embryos at the 1C, 2C, 4C and 8C stages were collected at 14-16 hours, 24 hours, 48 hours, and 72 hours, respectively. Embryo images were acquired using a Zeiss LSM 880 confocal microscope (Carl Zeiss AG, Germany). Mouse oocytes injected with various sperm by ICSI were transferred to KSOM medium and cultured at 37°C until collection. Embryos were collected at the following time post-ICSI: zygotes at 12 hours, E2C embryos at 20 hours, M2C embryos at 28 hours, L2C embryos at 34 hours, 4C embryos at 48 hours, and 8C embryos at 52 hours.

### In vitro transcription and microinjection

For mRNA preparation, the T7 promoter sequence and poly(A) tails of 100 adenines were added to the genes by PCR amplification. Gel-purified PCR products were used as templates for in vitro transcription with the mMESSAGE mMACHINE® Kit (Invitrogen, catalog no. AM1344) according to the manufacturer’s instructions. To knock down *Eif1a-like* genes, 5-10 picoliters of 20 μM antisense oligonucleotides synthesized by Hippobiotec were injected into fertilized zygotes. All injections were performed using an Eppendorf microinjector. For rescue experiments following *Eif1a-like* knockdown, 5-10 pl of 1.25 ng/μl codon-optimized *Eif1ad5-GFP* (ASO-resistant) mRNA together with 20 μM antisense oligonucleotides (ASOs) were injected. To visualize Eif1ad5-GFP localization in 2C embryos, 5-10 pl of 100 ng/μl mRNA was injected.

### Cell culture

The inducible 2CLC reporter line (*2C::tdTomato*; *Tet-on::Dux*) was obtained from B. Zhu (Institute of Biophysics, Chinese Academy of Sciences, Beijing, China)(*1*). mESCs were maintained under feeder-free conditions on tissue-culture plastic coated with 0.2% gelatin (Sigma-Aldrich, G1890) in N2B27 basal medium supplemented with 1 μM PD0325901, 3 μM CHIR99021 (2i), and 1,000 U/mL LIF. Cells were cultured at 37 °C with 5% CO₂, with daily medium changes, and passaged every 2–3 days using Accutase at a splitting ratio of 1:10 to 1:20. Cells were routinely tested for mycoplasma contamination

### Generation of *Eif1a-like* cluster knockout cells

The *Eif1a-like* cluster was deleted in mESCs using a dual-sgRNA CRISPR–Cas9 strategy. Four sgRNAs targeting the 5′ and 3′ flanking regions of the cluster (two per boundary) were cloned into the AAV-hU6-sgRNA-mU6-sgRNA vector backbone (plasmids QD187 and QD193). The sgRNA vectors were co-transfected with the Cas9-expressing plasmid pX330 (Addgene, 42230) into mESCs using JetPrime transfection reagent (Polyplus-transfection, 101000046) following the manufacturer’s protocol. At 48 h post-transfection, cells were subjected to single-cell cloning by limiting dilution. Genomic DNA from individual clones was screened by PCR using primers spanning the predicted deletion junction (F1/R2, 608 bp) and primers internal to the cluster (F1/R1, 999 bp; F2/R2, 1374 bp) to confirm biallelic deletion.

### Immunostaining

For mouse embryos, samples were fixed in 4% paraformaldehyde (G-CLONE, catalog no. PN4204) for 30 minutes at room temperature and then permeabilized with 0.5% Triton X-100 in PBS for 30 minutes. After blocking in 1% BSA in PBS for 1 hour, the embryos were incubated with the indicated primary antibody (β-Tubulin, A12289, ABclonal, 1:200) overnight at 4°C and then incubated with the appropriate fluorophore-conjugated secondary antibodies for 1-2 hours at room temperature in the dark. Nuclei were counterstained with DAPI (Life Technologies, D3571, 25 μg/ml) for 20 minutes. The embryos were then washed twice in 0.1% Triton X-100 in 1% BSA/PBS for 5 minutes each and mounted in antifade mounting medium. All immunofluorescence images were acquired using a Zeiss LSM700 confocal microscope.

For porcine and bovine embryos, fixation and permeabilization were carried out simultaneously in 1% paraformaldehyde and 0.2% Triton X-100 diluted in Dulbecco’s phosphate-buffered saline (DPBS), followed by overnight incubation at 4 C. After fixation, samples were washed in DPBS containing 0.001% PVA and blocked with 1% BSA for 1 hour at room temperature. Primary antibodies were diluted in blocking buffer and incubated with the samples overnight at 4 C (EIF1A, Abcam, ab172623, 1:100). After extensive washing in DPBS containing 0.001% PVA, samples were incubated with the appropriate fluorophore-conjugated secondary antibodies for 1-2 hours at room temperature in the dark. Nuclei were counterstained with DAPI for 20 minutes. Images were acquired using a Leica SP8 confocal microscope, and raw images were projected along the z-axis.

### Electrophoretic mobility shift assays

Recombinant EIF1AD8 was expressed in E. coli BL21DE3 as an MBP-fusion protein and purified; the MBP tag was removed, and purity was verified by SDS–PAGE. 601 Double-stranded DNA probes were prepared by annealing complementary oligonucleotides. Binding reactions contained 50 nM dsDNA and increasing concentrations of EIF1AD8 (0–3.2 μM) in reaction buffer (25 mM HEPES pH7.6, 1mM EDTA, 1mM TCEP). Reactions were incubated at 4 °C for 30 mins and resolved on 6% native polyacrylamide gels pre-run in 0.5× TBE at 120 V for 90 mins.

### RNA-seq library preparation and data processing

Oocytes and embryos were washed three times in medium containing 0.1% BSA in PBS and then lysed in 2 ul lysis buffer supplemented with RNase inhibitor. All RNA-seq libraries were generated using either the Smart-seq2 or Smart-seq3 protocol(*2, 3*). Paired-end RNA-seq reads were quality trimmed and aligned to the corresponding genomes using HISAT2 v2.2.1: mm10 for mouse, susScr11 for pig, and bosTau9 for bovine samples(*4*). Cufflinks v2.2.1(*5*)was used to estimate fragments per kilobase of transcript per million mapped reads (FPKM) for each gene based on Ensembl IDs from the Ensembl genome annotation database. FeatureCounts v2.0.3(*6*) was used to determine gene-level read counts with default settings. For hybrid embryo RNA-seq analysis, paired-end reads were classified into three categories: reads derived exclusively from the maternal species, reads derived exclusively from the paternal species, and reads potentially originating from both parents. Subsequent gene quantification and analysis were performed based on these classifications. To characterize dynamic gene expression patterns, heatmaps of stage-specific expression profiles were generated using R package pheatmap v1.0.12. For visualization, extreme FPKM values were capped at 0.001 and 0.64 quantiles. Differentially expressed genes (DEGs) were identified using DESeq2 v1.44.0(*7*) with adjusted P < 0.05 and FC > 2. Gene Ontology analysis of DEGs was performed using clusterProfiler v4.10.1(*8*).

### Identification of ZGA genes and stage-specific genes

ZGA genes and stage-specific genes were defined based on the reference RNA-seq data(*9*). Using mouse embryonic data as an example, ZGA genes were defined as genes that were not expressed or were lowly expressed in full-grown and MII oocytes (FPKM < 1) but were upregulated (FPKM > 5) in either 1C or E2C embryos (minor ZGA genes) or in L2C embryos (major ZGA genes). Genes specifically activated at each stage during early development were defined using stricter criteria: expression at the defined stage (FPKM > 5) but silencing in the immediately preceding stages (FPKM < 1). In bovine and porcine embryos, ZGA genes were identified as genes that were not expressed or only minimally expressed in GV and MII oocytes (combined GV + MII FPKM < 4) but showed marked upregulation in zygote, 2C, 4C, or 8C embryos (FPKM > 5). Genes specifically activated at one stage during early development were defined using the stricter criteria described above (FPKM > 5 at that stage and FPKM < 3 in the immediately preceding stages). In hybrid embryos, stage-specific genes were defined in a corresponding manner, with genes expressed in inbred embryos (FPKM > 5 at any stage) removed from the list of stage-specific genes in hybrid embryos.

### Identification of conserved ZGA genes

To identify conserved ZGA genes in the mouse genome, we compared stage-specific genes from inbred mouse embryos at the 2C, 4C, and 8C stages with those from hybrid mouse embryos after the 2C stage. Genes shared by both groups were defined as conserved ZGA genes in mice. We then performed Gene Ontology (GO) analysis using clusterProfiler v4.10.1 and grouped these genes into functional clusters(*10*).

### Identification of housekeeping genes

For housekeeping gene identification, embryonic samples were supplemented with RNA-seq data from multiple organs of the same species, including heart, liver, spleen, lung, kidney, and others. Genes showing stable expression (FPKM > 10) across all these samples were defined as housekeeping genes(*11*).

### ChIP-seq library preparation

ChIP-seq was performed as previously described(*12*) with minor modifications. Briefly, cells were dissociated with Trypsin-EDTA, washed once in PBS, and cross-linked in growth medium containing 1% formaldehyde for 10 min at room temperature with gentle rotation. Cross-linking was quenched by adding glycine to a final concentration of 0.125 M for 5 min at room temperature. Cells were washed twice with ice-cold PBS, lysed in nuclear lysis buffer on ice for 10 min, and nuclei were collected by centrifugation. Chromatin was fragmented to 200–500 bp by MNase digestion followed by sonication using a Covaris ultrasonicator.

Soluble chromatin was pre-cleared for 2 hours with 50 μl of a 1:1 mixture of Protein A and Protein G Dynabeads (Thermo Fisher Scientific, 10002D/10004D) pre-bound with 3 μg rabbit or mouse IgG. The indicated antibody (GFP, Abcam, ab290) was then incubated with the lysates overnight at 4°C with rotation, after which 50 μl Dynabeads were added and incubation continued for an additional 2 hours. After five washes with high-salt wash buffer and a final wash with TE buffer, the beads were resuspended in 100 μl TE elution buffer and digested with RNase Cocktail (Thermo Fisher Scientific, AM2286) for 30 minutes at 37°C, followed by proteinase K (1 mg/ml) for 1 hour at 42°C. The samples were then incubated overnight at 65°C and purified using a QIAquick PCR purification kit (Qiagen, 28104). Sequencing libraries were prepared as previously described.

### CUT&Tag library preparation

CUT&Tag assays were performed as previously described(*13*) with minor modifications. Briefly, cells were mildly cross-linked in 1 mL PBS containing 0.1% formaldehyde for 2 min at room temperature with gentle rotation, and cross-linking was quenched by adding glycine to a final concentration of 125 mM for 5 min at room temperature. Cells were washed once with 1 mL wash buffer with gentle pipetting and collected by centrifugation. Concanavalin A-coated magnetic beads (Bangs Laboratories, BP531) were activated in 100 μl binding buffer per 10 μl bead slurry and incubated with 2 × 10⁵ cells for 10 min at room temperature. Bead-bound cells were washed once in antibody buffer and resuspended in 50 μl antibody buffer containing the indicated primary antibody (GFP, Abcam, ab290; EIF1A, ab20954; EIF1A, ab172623). The mixture was rotated overnight at 4 °C, followed by incubation with the secondary antibody (guinea pig anti-rabbit IgG H&L, ABIN101961) for 1 h at room temperature. After three washes with 500 μl dig-wash buffer, cells were resuspended in 100 μl dig-300 buffer and incubated with the pAG-Tn5 adapter complex while rotating at 300 rpm for 1 h at 25 °C. Following three washes with dig-300 buffer, cells were resuspended in 300 μl Tagmentation buffer and incubated at 37 °C for 1 h. Tagmentation was terminated by adding 10 μl of 0.5 M EDTA (pH 8.0), 3 μl of 10% SDS, and 2.5 μl of proteinase K (20 mg/mL), followed by incubation at 50 °C for 1 h. The products were purified and PCR amplified as described for sequencing on an Illumina NovaSeq 6000 platform.

### ATAC library preparation

ATAC-seq libraries were prepared as previously described with minor modifications(*9*). Briefly, samples were lysed in 11 μl lysis buffer (10 mM Tris-HCl, pH 7.5, 10 mM NaCl, 3 mM MgCl_2_, and 0.05% digitonin) for 10 minutes at 4°C. Samples were then incubated with 5 μl Tn5 transposase and 4 μl TTBL buffer at 37°C for 30 minutes (Vazyme, TD502). Following tagmentation, 2 μl 10% SDS was added directly to terminate the reaction. Next, 2 μl each of carrier RNA and spike-in DNA were added, and PCR was performed for 14 cycles to amplify the library. Libraries were then purified by 0.4x-2x AMPure bead size selection and subjected to next-generation sequencing.

### ChIP-seq, CUT&Tag and ATAC-seq data processing

Adapters from ChIP-seq, CUT&Tag and ATAC-seq data were removed using cutadapt v4.5. The trimmed reads were then processed with bowtie2 v2.5.2(*14*) to remove mitochondrial reads, followed by alignment to the mm10 genome with bowtie2 v2.5.2 using the following parameters: -t -L 25 -X 2000 --no-mixed --no-discordant. Unmapped reads, non-uniquely mapped reads, and PCR duplicates were removed. Genome-wide read coverage over mm10 was calculated with bamCoverage from deepTools v3.5.4(*15*) using the parameters --binSize 50 --smoothLength 150 --skipNAs --normalizeUsing RPKM and visualized with IGV v2.19.2(*16*). Peaks were called by MACS v2.2.9.1(*17*) using the parameters -f BAMPE -q 0.05 --keep-dup all --nomodel. The numbers of overlapping peaks were calculated using BEDTools v2.31.0(*18*).

### Motif-enrichment analysis

The resulting peaks were classified into genic and distal regions using ChIPseeker(*19*), and peak summit sites were extracted simultaneously. Motif analysis was then performed on the different regions using HOMER(*20*) with the parameters -p 20 -S 12,10,8 -size −20,20. Motif enrichment within peaks was calculated using FIMO v5.5.8(*21, 22*) with the parameter --thresh 1e-3.

### Co-immunoprecipitation and mass-spectrometry analysis

Co-immunoprecipitation and mass spectrometry were performed as previously described with minor modifications(*23*). Briefly, E5-GFP proteins from 2CLC or mESC samples were individually precipitated with GFP-Trap MA bead slurry (Allele Biotech). The samples were rotated overnight at 4°C and washed seven times with buffer containing 10 mM Tris-HCl (pH 7.4), 500 mM NaCl, 0.1% Triton X-100, and 0.1 mM PMSF. Proteins bound to the beads were boiled in sample buffer and analyzed by one-dimensional gel electrophoresis. The gel was excised, sliced, and subjected to in-gel trypsin digestion. Mass spectrometric analysis of the purified samples was performed by liquid chromatography-tandem mass spectrometry (LC-MS/MS) using a Linear Ion Trap mass spectrometer (Thermo Fisher Scientific) equipped with a nanospray source and Eksigent high-performance liquid chromatography. Peak areas of fragment ions were used to calculate the relative intensities of precursor ions for selected peptides. At least two peptides were used to quantify each protein. The mean relative intensity of the selected peptides represented the relative expression level of each protein. Average enrichment between bait and control samples was calculated using a custom R script, and adjusted P values were calculated using the R package limma(*24*).

## Data availability

All data is available within the article and its Supplementary Tables. All newly generated data have been deposited in GEO under accession numbers GSE328869 (RNA-seq), GSE328840 (ATAC-seq), GSE328839 (ChIP-seq and CUT&Tag). The mass spectrometry proteomics data have been deposited to the ProteomeXchange Consortium via the PRIDE partner repository with the dataset identifier PXD078112. Publicly available datasets used in this study include *Obox* KO RNA-seq data from GSE215813(*9*), *Obox* OE RNA-seq data from GSE196671(*25*), *Dux* OE RNA-seq data from GSE85627(*26*), 2CLC and mESC RNA-seq data from GSE75751(*27*), and 2CLC H3K27ac/H3K4me3/H3K4me1 CUT&Tag data from GSE159621(*28*).

## Code availability

The software used to analyze these data is listed in the Methods and is publicly available. All related code is available on GitHub (https://github.com/BioInfrookie/Mammalian_ZGA).

